# Selective interaction of the protein SMCHD1 with specific chromatin regions is governed by the loading factor LRIF1 and SMCHD1 ATPase activity

**DOI:** 10.1101/2025.06.13.659515

**Authors:** Flavia Constantinescu, Aleksander T Szczurek, Tatyana B Nesterova, Guifeng Wei, Victoria Pustygina, Stephan Uphoff, Adam D Cawte, Neil Brockdorff

## Abstract

The chromosomal protein SMCHD1 is a GHKL ATPase that plays important roles in epigenetic silencing, including on the inactive X chromosome (Xi) and at the D4Z4 macrosatellite linked to regulation of DUX4 expression in the disorders facioscapulohumeral muscular dystrophy (FSHD2) and Bosma arrhinia micropthalmia syndrome (BAMS). In this study we use live-cell and single-molecule imaging approaches to investigate SMCHD1 interactions with chromatin and its function in epigenetic silencing. We show that chromatin binding of SMCHD1 genome-wide, including on the Xi, is critically dependent on the protein LRIF1 that mediates interaction with H3K9me2/3 modified nucleosomes. Using engineered mutations in the GHKL ATPase domain we show that ATP hydrolysis is required for selective enrichment of SMCHD1 at specific chromatin regions, which is critical for gene silencing on the Xi. A gain-of-function mutation, G137E, that occurs in BAMS patients, results in accelerated Xi recruitment and greater Xi chromosome compaction. Together, our findings advance mechanistic understanding of SMCHD1 function on the Xi and at other target sites in the genome.

## Introduction

X-chromosome inactivation (XCI) evolved in mammals to ensure dosage compensation of X-linked genes in XX females relative to XY males^1^. The master regulator of this process is the X-inactive specific transcript (Xist), a 15-18 kb long noncoding RNA (lncRNA) that accumulates *in cis* over the future inactive X (Xi) chromosome^2–5^. Xist mediates stepwise recruitment of factors which modify the underlying chromatin and ultimately lead to chromosome-wide silencing that is then maintained through all subsequent cell divisions. One of these factors is the epigenetic modifier structural maintenance of chromosome hinge domain containing 1 (SMCHD1), whose absence causes female-specific mid-gestation embryonic lethality ^6,7^. SMCHD1 has two characteristic conserved domains, a C-terminal SMC hinge domain, found also in condensin, cohesin, and other proteins belonging to the SMC family of chromosomal proteins^8^, and an N-terminal GHKL ATPase domain, also found in a range of ATPases including DNA gyrase and the MORC family of epigenetic repressors^9^.

SMCHD1 is recruited to Xi at a relatively late stage following initiation of XCI^10,11^, leading to suggestions that it is principally involved in stable maintenance of the inactive state. However, more recent work has determined that SMCHD1 contributes to the silencing of a significant subset of X-linked genes, referred to as SMCHD1-dependent and SMCHD1-partially dependent genes respectively^11–14^. Additionally, SMCHD1 function is required for DNA methylation of CpG-islands of most X-linked genes, a hallmark of stable long-term silencing, and for establishing the unique long-range 3D architecture of the Xi^10,13–17^. Finally, there is evidence suggesting that SMCHD1 contributes to the compacted structure of Xi chromatin^18^. The mechanism for SMCHD1 recruitment to Xi is not fully understood, although it is known to require the histone modification H2AK119Ub catalysed by the Polycomb repressive complex PRC1^19^.

In addition to its role in XCI, SMCHD1 functions in the epigenetic repression of certain autosomal loci^15,16,20,21^. A notable case is the gene encoding DUX4 in human, abnormal expression of which results in FSHD2. Here SMCHD1 plays a role in DUX4 repression via the D4Z4 macrosatellite array, and loss-of-function mutations in SMCHD1 are causative in a proportion of FSHD2 patients^22–24^. FSHD2 mutations have also been found in a second factor, LRIF1, which together with proteins of the HP1 family, directs SMCHD1 to target sites marked by the histone modification H3K9me3^18,25,26^. Abnormal expression of DUX4 also occurs in the rare disorder BAMS, and this has been linked to SMCHD1 gain-of-function mutations in the GHKL ATPase domain^23,24,27,28^.

Structural analysis of SMCHD1 has shown that it forms a stable homodimer with an extended rod-like structure^26^. Dimerisation is mediated by the SMC hinge and the GHKL ATPase domains^26,29–31^. The functional importance of the GHKL ATPase is underscored by its requirement for SMCHD1 association with the Xi^26,32^, and the occurrence of GHKL ATPase mutations in FSHD2 and BAMS patients. However, the role of the GHKL ATPase in chromosomal targeting and/or function of SMCHD1 in epigenetic repression is not well understood.

Here we employ live-cell imaging and single-particle tracking (SPT) to study the dynamic behaviour of SMCHD1 and, thus, shed light on the molecular mechanisms underlying its interaction with chromatin and functions in XCI. We show that LRIF1 plays an important role in loading SMCHD1 at target sites including pericentric heterochromatin (PCH) and Xi. Furthermore, we demonstrate that SMCHD1 exhibits Xi-specific dynamic behaviour which is altered by mutations affecting the GHKL ATPase activity of the protein, including loss- and gain-of-function mutations associated with BAMS and FSHD2. Importantly, these mutations also affect SMCHD1 function in X-linked gene silencing and chromosome compaction.

## Results

### An mESC model for analysing SMCHD1 dynamic behaviour

To investigate SMCHD1 association with Xi we adapted iXist-ChrX, a previously established interspecific XX mESC line with the TetOn promoter driving doxycyline inducible Xist RNA expression on the 129S strain-derived X chromosome^33^. We first engineered a Halo-tag^34^ on the native SMCHD1 protein (Figure 1a, b). Prior experiments (using a SNAP-tag^35^), indicated that C-terminal tagging affects SMCHD1 stability (Extended Data Figure 1a), but an N-terminal Halo-tag had no detectable effect on the protein level (Figure 1b). Furthermore, allelic transcription analysis using ChrRNA-seq^33^ affirmed that addition of the N-terminal Halo-tag had no effect on SMCHD1 function in XCI (Extended Data Figure 1b).

**Figure 1.**
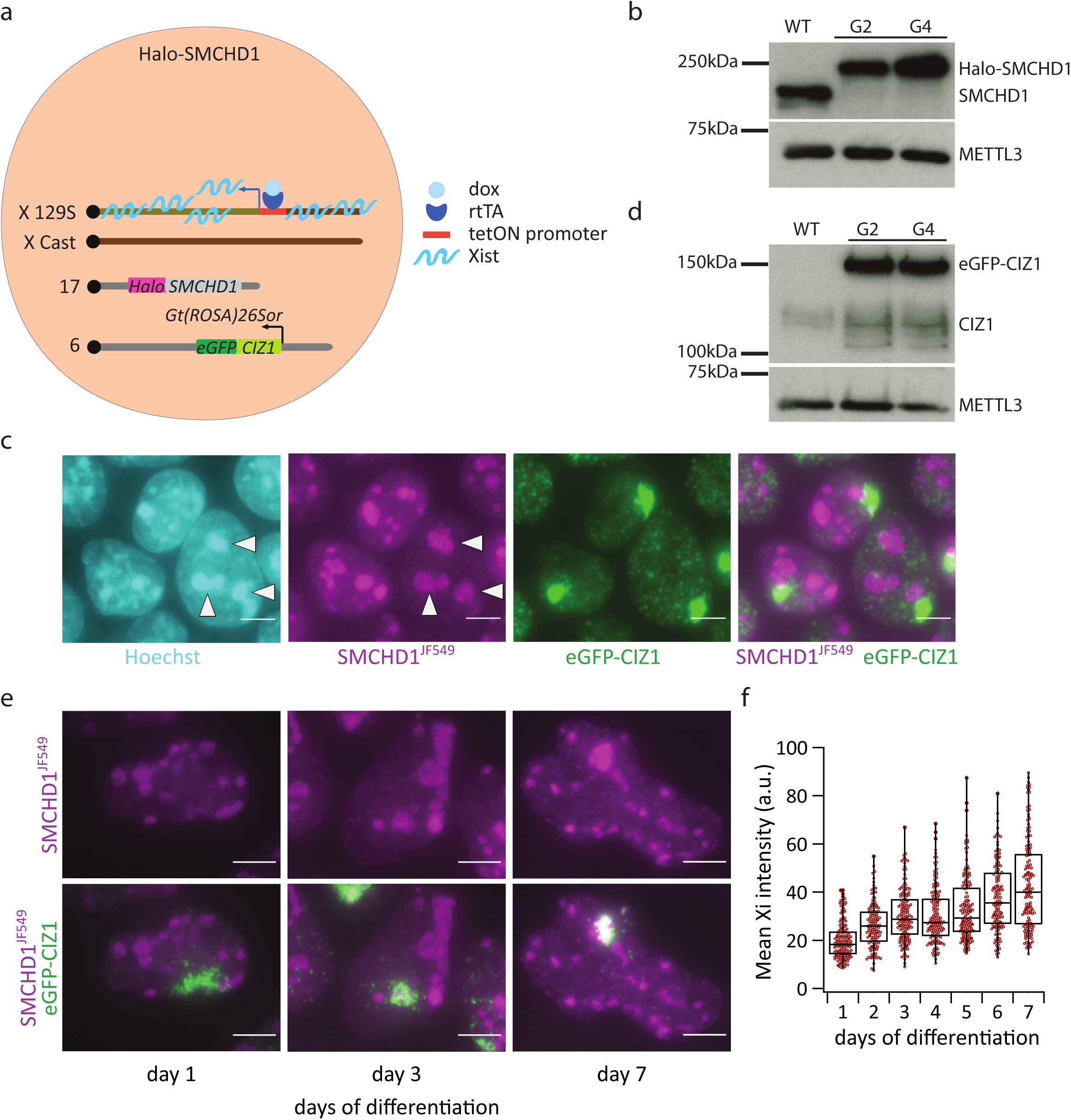
A model system to analyse dynamic interactions of SMCHD1 with chromatin. a. Schematic of the Halo-SMCHD1 XX mESC line generated in the iXist-ChrX mESC line. b. Western Blot showing the level of expression of Halo-SMCHD1 in two independent clones. Clone G2 was used for further experiments. METTL3 loading control c. Representative live-cell images of Halo-SMCHD1 mESC line, taken after 24h doxycycline addition. Arrowheads indicate selected PCH domains. Images are maximum-intensity projections, scale bar - 5µm. d. Western blot showing the expression of eGFP-tagged CIZ1 in two independent clones. METTL3 loading control e. Representative live-cell images showing gradual accumulation of SMCHD1 over the Xi domain marked by eGFP-CIZ1 during mESC differentiation. Images are maximum-intensity projections, scale bar - 5µm. f. Quantification of SMCHD1 signal underlying the CIZ1 domain at each day during a 7-day live-cell imaging differentiation timecourse. Each dot represents a single cell. n = 136-159.

Labelling of endogenously tagged SMCHD1 with the fluorescent Halo ligand JF549 revealed nuclear localized protein as expected. Interestingly, the majority of cells exhibited enrichment of SMCHD1 at pericentric heterochromatin (PCH) regions as determined by staining with Hoechst 33342 (Figure 1c), contrasting with our previous analysis of SMCHD1 localisation by immunofluorescence (IF) in fixed mESCs^26^. Further investigation of this finding is described below.

To identify the Xi domain in live-cell imaging we introduced a transgene encoding eGFP-CIZ1 in the *Gt(ROSA)26Sor* locus (Figure 1a). Expression of eGFP-CIZ1 was confirmed by western blot (Figure 1d). CIZ1 binds to the E-repeat element of Xist RNA from the onset of XCI and continuously thereafter^36,37^, and consistent with this, eGFP-CIZ1 Xi domains were detected in the majority of cells following Xist RNA induction in undifferentiated mESCs (Figure 1c, Extended Data Figure 1c). RNA-FISH analysis in mESCs confirmed the colocalization of eGFP-CIZ1 and Xist RNA (Extended Data Figure 1d).

In previous work using IF in fixed cells we observed enrichment of SMCHD1 over the Xi only after several days of Xist RNA induction coupled to cellular differentiation^11^. To investigate Xi localisation of SMCHD1 by live-cell imaging we used a monolayer differentiation protocol^38^. We then quantified Halo-SMCHD1 enrichment over the Xi (eGFP-CIZ1 domains) in a live-cell imaging timecourse over 7 days of differentiation (Figure 1e, f). Strong accumulation of SMCHD1 over the Xi was most clearly evident at late differentiation stages (Figure 1e), consistent with prior studies^10,11^. However, quantitative analysis demonstrated a more graded accumulation of SMCHD1, with significant enrichment underlying CIZ1 domains detectable as early as 48 hours after Xist RNA induction (Figure 1f).

### SMCHD1 localisation to PCH is LRIF1 dependent

Prior analysis of SMCHD1 in mESCs using IF revealed enrichment at PCH regions in only a small proportion of cells^26^. Similarly, IF in the mESC line analysed herein does not show PCH accumulation of SMCHD1 in most cells (Extended Data Figure 2a). However, Halo-tag labelling followed by formaldehyde fixation displays the same population-wide pattern observed in live-cell imaging (Extended Data Figure 2b), suggesting that absence of this localisation pattern using IF is not attributable to fixation but likely due to epitope masking when the protein is bound to PCH regions. We note that these observations support findings from a proteomic analysis that defined SMCHD1 amongst proteins enriched at PCH in mESCs^39^.

PCH accumulation of SMCHD1 could be explained by its reported interaction with LRIF1 and HP1 family proteins to mediate association with the chromatin mark H3K9me3^18,26^. To explore this hypothesis, we used CRISPR/Cas9 genome engineering to generate iXist-ChrX mESCs with a Halo-tag on the C-terminus of LRIF1 (tagging all three known isoforms) (Extended Data Figure 2c), and then performed live-cell imaging after labelling cells with fluorescent JF549 Halo ligand. As shown in Figure 2a, LRIF1-Halo is strongly enriched over PCH domains in a pattern that is indistinguishable from that seen for Halo-SMCHD1. Localisation of LRIF1 to PCH domains has been reported previously^40^, but was seen in only a small proportion of cells in a population, probably due to the use of antibody-based staining rather than live-cell imaging.

**Figure 2.**
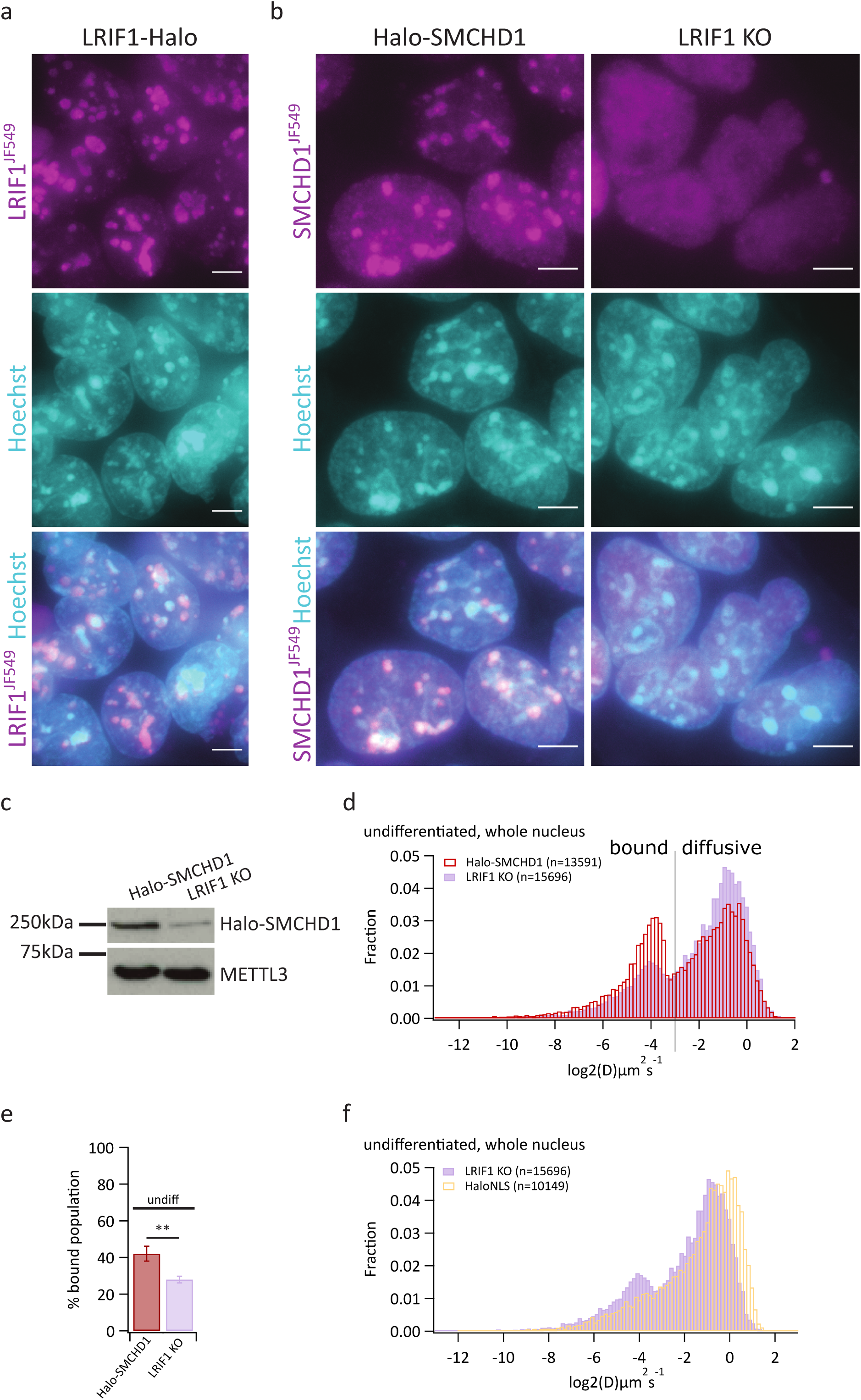
LRIF1 is required for SMCHD1 binding to chromatin in mESCs. a. Representative live-cell images of LRIF1-Halo mESC line, taken 24h after doxycycline addition. b. Representative live images of Halo-SMCHD1 and Lrif1^-/-^ (LRIF KO) mESC lines. c. Western blot showing SMCHD1 protein levels in the presence and absence of LRIF1. d. Normalized histogram of diffusion coefficients of tracks from across the whole nucleus. Cells were undifferentiated, kept for 24h with doxycycline. e. Bar plots showing the average percentage of bound molecules of SMCHD1 (3 independent replicates each). Error bars represent the STDEV. Statistical analysis was by unpaired t-test. f. In purple, same as in 2d. In yellow, normalized histogram of diffusion coefficients of tracks from across the whole nucleus; cells are undifferentiated Halo-3xNLS mESCs, kept for 24h with doxycycline. All images are maximum-intensity projections. Scale bars correspond to 5µm. undiff = undifferentiated

To test if LRIF1 is required for SMCHD1 localisation at PCH domains we generated CRISPR/Cas9-mediated knock-out of the gene encoding LRIF1 in the Halo-SMCHD1 mESC line described above (Extended Data Figure 2c-e). The engineered gene knock-out was designed to ensure loss of all three LRIF1 isoforms. The *Lrif1^-/-^* mESCs exhibit a marked loss of SMCHD1 enrichment at PCH regions in most cells, even though the regions are still clearly present as highlighted by staining with Hoechst 33342 dye (Figure 2b). This observation confirms that LRIF1 plays a central role in the observed recruitment of SMCHD1 to PCH. SMCHD1 localisation to PCH regions remain detectable in a small proportion of cells (Extended Data Figure 2f). Of note, absence of LRIF1 also leads to a dramatic decrease in the protein level of SMCHD1 (Figure 2c).

To assess changes in the dynamic behaviour of SMCHD1, we made use of a photoactivable Halo-tag ligand (PA-JF549), and highly inclined and laminated optical sheet (HILO) microscopy with high temporal resolution (15ms frames)^41^. This allowed us to perform Single Particle Tracking (SPT) analysis to record the localisations, movement, and diffusion coefficients of individual Halo-SMCHD1 molecules in live cells. In wild-type cells, a histogram plot of the diffusion coefficients shows two populations, an immobile, chromatin bound population with a Log_2_ diffusion coefficient peak around -4 (0.0625 µm^2^/s) and a fast, freely diffusing, population with a peak at around -1 (0.5 µm^2^/s) (Figure 2d). The bound population is defined herein as the percentage of tracks whose Log2 diffusion coefficient is smaller than -3 (Figure 2e). The threshold value -3 was chosen as it sits, across the various conditions analysed in this study, close to the minimum between the peaks of bound and unbound populations. Importantly, this analysis method captures differences between biological contexts with a similar efficiency as the methods relying on Gaussian fitting or kinetic modelling in SPOT-ON, but does not require large data-sets^42,43^ (Extended Data Figure 2g).

In *Lrif1^-/-^* mESCs, the bound population is significantly reduced (Figure 2d-e), exhibiting similar dynamic behaviour to Halo-tag protein with a nuclear localization signal (Halo-3xNLS) which is not expected to interact with chromatin (Figure 2f). These observations are consistent with subcellular fractionation experiments which reported loss of SMCHD1 from the chromatin bound fraction in cells lacking LRIF1^26^. We speculate that loss of chromatin binding increases SMCHD1 protein turnover, accounting for reduced levels of the protein in *Lrif1^-/-^* mESCs (Figure 2c).

### SMCHD1 exhibits enhanced binding over the Xi territory

We went on to analyse the dynamic behaviour of SMCHD1 in the context of XCI and cellular differentiation. We established an analysis pipeline to define SMCHD1 diffusion behaviour underlying the eGFP-CIZ1-marked location of the Xi domain (Extended Data Figure 3a, b). As a control, for each analysed Xi territory we used a randomly selected, equally-sized nuclear region (Extended Data Figure 3b). Using this approach we observed Xi-specific dynamic behaviour of SMCHD1 in both undifferentiated (Figure 3a, c) and differentiated (Figure 3b, c) mESCs. In both cases there is a more prominent bound population over the Xi compared to random regions, where SMCHD1 dynamics are, as expected, similar to those observed nucleus-wide (Figure 2d). Notably, the profile of SMCHD1 diffusion coefficients over the Xi region in differentiated cells strongly resembles that of Halo-tagged histone H2B (Extended Data Figure 3c). Increased binding of SMCHD1 over the Xi territory was confirmed by fluorescence recovery after photobleaching (FRAP) analysis, which revealed enhanced residency and immobile populations compared to the nucleoplasm (Extended Data Figure 3d-f).

**Figure 3.**
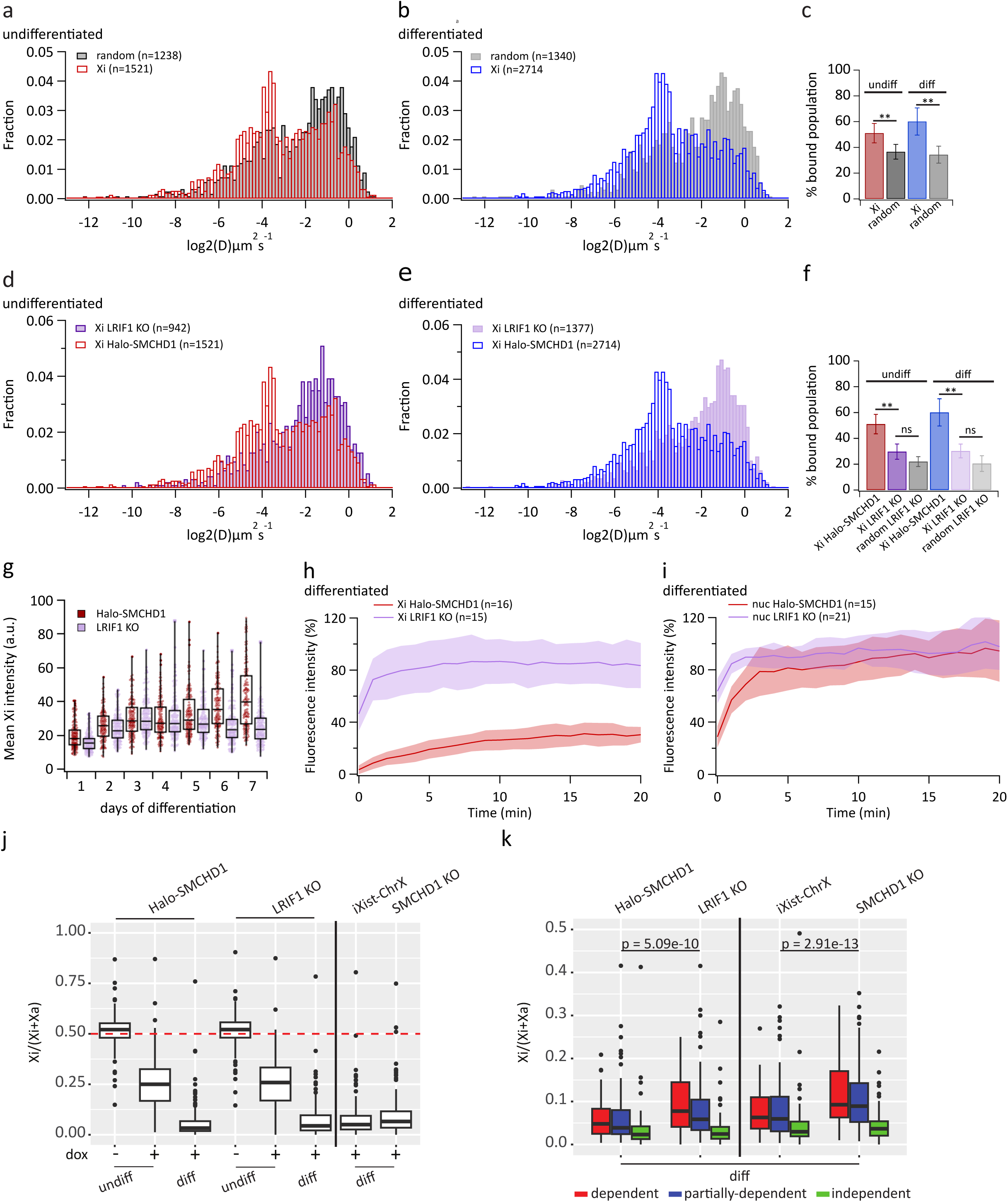
LRIF1 is required for SMCHD1 binding to Xi chromatin in differentiating mESCs. a. Normalized histogram of diffusion coefficients of tracks from Xi (red) or a random region (black) in undifferentiated Halo-SMCHD1 mESCs treated for 24h with doxycycline. b. Normalized histogram of diffusion coefficients of tracks from Xi (blue) or a random region in the nucleus (grey) in Halo-SMCHD1 mESCs differentiated for 7 days. c. Bar plot showing the average percentage of bound molecules of SMCHD1 (5 independent replicates each). Error bars represent the STDEV (unpaired t-test). d. In red, same as in 2a. In dark purple, normalized histogram of diffusion coefficients of tracks from Xi in undifferentiated Lrif1^-/-^ mESCs treated 24h with doxycycline. e. In blue, same as in 2b. In light purple, normalized histogram of diffusion coefficients of tracks from Xi in Lrif1^-/-^ mESCs differentiated for 7 days. f. Bar plot showing the average percentage of bound molecules of SMCHD1 (5 or 4 independent replicates for Halo-SMCHD1 and Lrif1^-/-^ mESCs, respectively). Error bars represent the STDEV (unpaired t-test). g. Quantification of SMCHD1 signal underlying CIZ1 domains during a 7-day differentiation timecourse comparing Halo-SMCHD1 (red) and Lrif1^-/-^ (purple) mESCs. Each dot represents a cell. n = 136-200. h-i. Average fluorescence recovery curves over Xi (h) or a random nucleoplasmic region (i) comparing Halo-SMCHD1 and Lrif1^-/-^ mESCs differentiated for 7 days. Shading represents the STDEV. j. Box plot showing allelic X-linked gene expression in indicated mESCs with and without doxycycline (dox) and 7 day differentiation. iXist-ChrX is the matched parental line for SMCHD1 KO mESCs. Samples represent one or an average of two replicates. k. Box plot showing allelic chrRNA-seq analysis of transcription of X-linked genes in indicated cell lines differentiated for 7 days. Genes are split in three categories according to their dependency on SMCHD1 for silencing^13^. The samples represent an average of two or three replicates. Statistical analysis was paired t-test. undiff = undifferentiated, diff = differentiated

Previous studies found that LRIF1 depletion in fully differentiated XX somatic cells has no effect on SMCHD1 localisation to the Xi domain ^18,26^, although IF analysis demonstrated localisation of LRIF1 protein on the Xi. To revisit this issue using live-cell imaging we made use of the LRIF1-Halo line described above, which also includes eGFP-CIZ1 integrated into the *Gt(ROSA)26Sor* locus. This analysis indicated strong enrichment of LRIF1-Halo within the Xi territory in 7 day differentiated cells, although not at an earlier stage of 24h Xist induction (Extended Data Figure 4a,b). These observations prompted us to investigate if there is a role for LRIF1 in establishing SMCHD1 enrichment at the onset of XCI. Thus, we analysed Halo-SMCHD1 in the *Lrif1^-/-^*mESCs described above. We observed a strong reduction of bound SMCHD1 and a prominent fast diffusing population over the Xi, both in undifferentiated and differentiated cells (Figure 3d-f). This is accompanied by a clear reduction in Xi associated SMCHD1 (Figure 3g, Extended Data Figure 4c).

Importantly, the elevated bound population of SMCHD1 on the Xi is lost in the *Lrif1^-/-^* mESCs, with the population distributions within the Xi and a random region being similar, both in undifferentiated and differentiated mESCs (Extended Data Figure 4d-e, Figure 3f). These observations were corroborated by FRAP analysis in *Lrif1^-/-^*cells showing SMCHD1 has a similar recovery behaviour within the Xi territory and the nucleoplasm, with a faster and more complete recovery level compared to wild-type cells (Figure 3h-i, Extended Data Figure 4f-i). Consistent with these findings, we observed impaired X-linked gene silencing at day 7 of differentiation in *Lrif1^-/-^* cells, when normally X-linked gene repression is almost complete (Figure 3j). Impaired silencing was most evident for SMCHD1-dependent and partially dependent genes (as categorized in^13^), and is comparable to what is observed in *SmcHD1* KO mESCs (Figure 3k).

### SMCHD1 GHKL ATPase activity regulates chromatin enrichment

In previous work we reported that E147A mutation of the SMCHD1 GHKL ATPase domain abrogates Xi accumulation in XX somatic cells^26^. *In vitro* analysis has shown that E147A completely abolishes ATP hydrolysis, although ATP is still able to bind^26,30^. We set out to investigate how this mutation affects the dynamic behaviour of SMCHD1 on the Xi and genome wide. In parallel we tested a second mutation Y353C, identified in FSHD2 patients, that has been shown to strongly abrogate SMCHD1 ATPase activity without affecting folding/protein stability^24,27^. Using CRISPR/Cas9-mediated homologous recombination, we introduced each of these mutations, biallelically at the endogenous *SMCHD1* locus in the wild-type cell line, Halo-SMCHD1 (Extended Data Figure 5a,b). Western blot analysis indicated that both mutations result in a small reduction in levels of SMCHD1 (Extended Data Figure 5c,d). Both mutant proteins show loss of enrichment at PCH regions, as seen in *Lrif1^-/-^*cells (Figure 4a), and over the Xi during a 7-day differentiation time-course (Figure 4b, Extended Data Figure 5e,f). Consistent with these observations, ChrRNA-seq revealed that SMCHD1^E147A^ and SMCHD1^Y353C^ show abrogated Xi silencing (Figure 4c), particularly affecting SMCHD1-dependent genes and partially dependent genes (Figure 4d).

**Figure 4.**
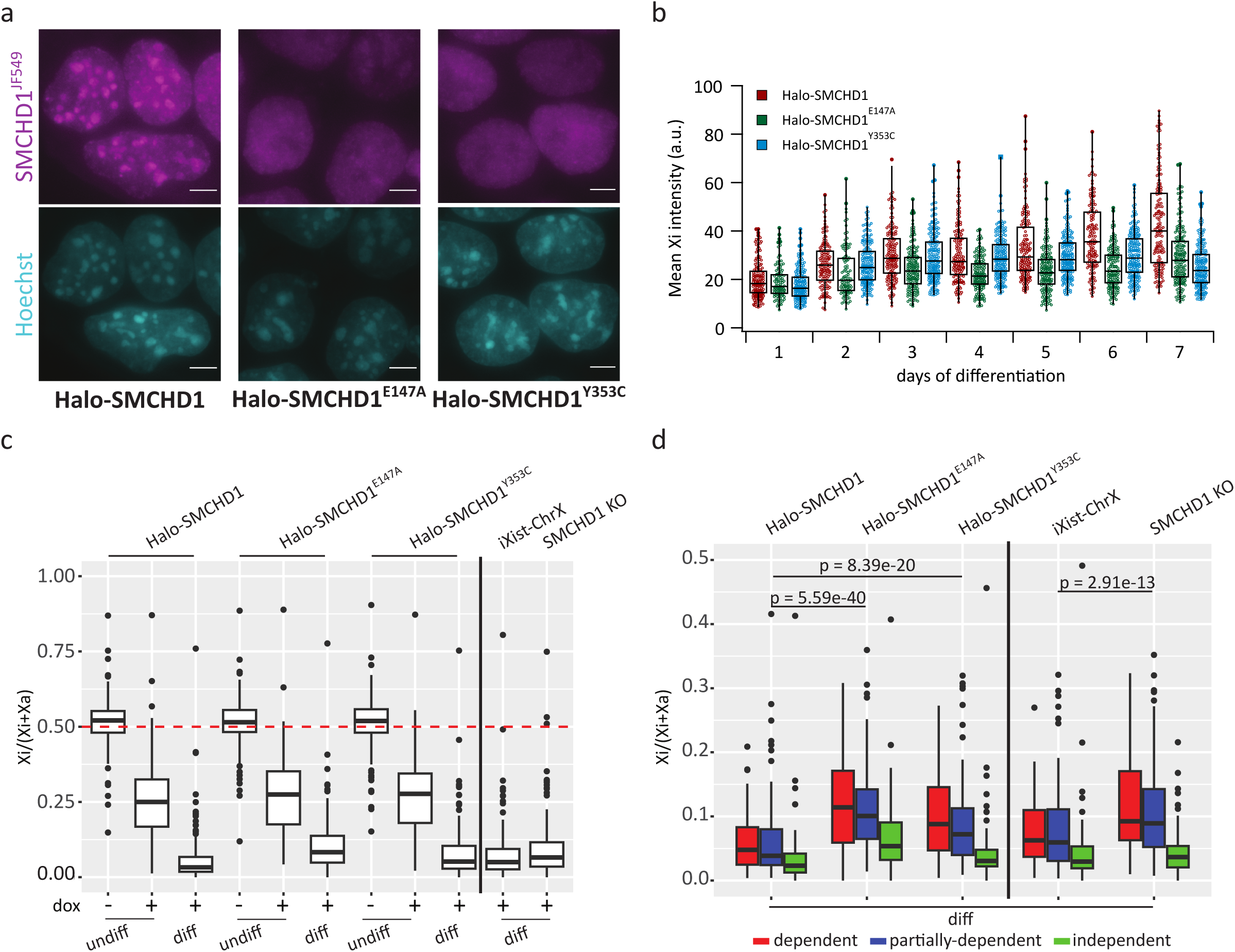
SMCHD1 GHKL ATPase mutations abrogate Xi localisation and silencing function. a. Representative live-cell images of indicated mESCs undifferentiated, kept for 24h with doxycyline. Images are maximum-intensity projections. Scale bar - 5µm. b. Quantification of SMCHD1 signal underlying CIZ1 domains in indicated cell lines during a 7-day differentiation timecourse. Each dot represents a single cell. n = 102-206. c. Box plot showing allelic ChrRNA-seq analysis of transcription of X-linked genes in indicated cell lines with and without doxycycline (dox) and 7 day differentiation. The boxes represent one or an average from two or three replicates d. Box plot showing allelic ChrRNA-seq analysis of transcription of X-linked genes in mESC lines which were differentiated for 7 days, with genes split in three categories according to their dependency on SMCHD1 for silencing^13^. An average of two or three replicates was used for this plot. Statistical analysis - paired t test. undiff = undifferentiated, diff = differentiated

Analysis of the dynamic behaviour of the mutant proteins across the whole nucleus showed that SMCHD1^E147A^ has a larger bound population compared to the wild-type protein, both in undifferentiated and differentiated cells (Figure 5a, b, Extended Data Figure 6a, b). SMCHD1^Y353C^ on the other hand has a distribution profile similar to wild-type (Figure 5c, d, Extended Data Figure 6c, d). These results highlight an interesting contrast between the *Lrif1^-/-^*, SMCHD1^E147A^ and SMCHD1^Y353C^ cells which all show loss of PCH accumulation of SMCHD1, whilst exhibiting different dynamic characteristics.

**Figure 5.**
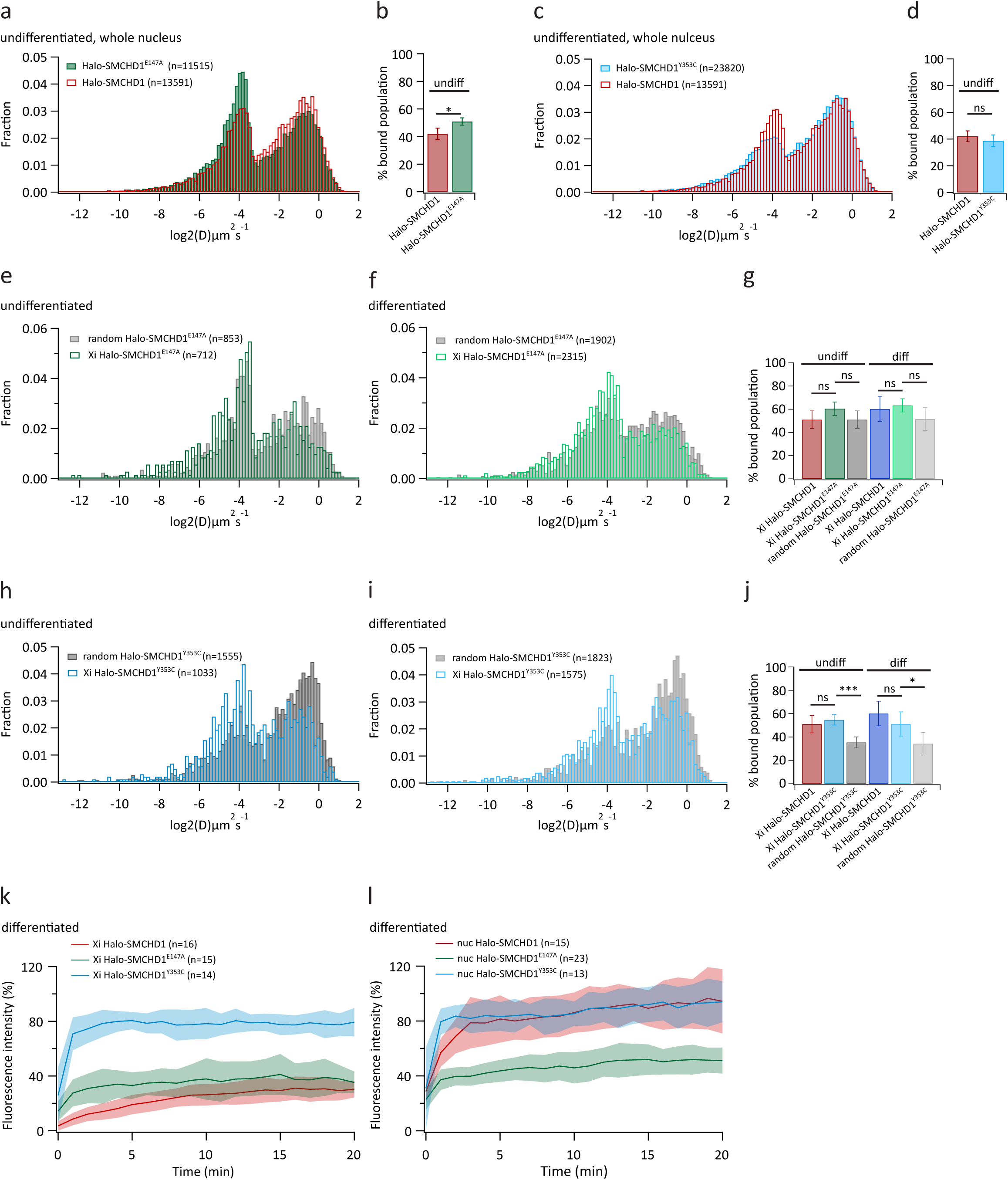
SMCHD1 GHKL ATPase loss-of-function mutations alter chromatin binding behaviour. a. In red, same as Figure 2d. In green, normalized histogram of diffusion coefficients of tracks from across the whole nucleus in undifferentiated Halo-SMCHD1^E147A^ mESCs, kept for 24h with doxycycline. b. Bar plots showing the average percentage of bound molecules of SMCHD1 (3 independent replicates each). Error bars represent the STDEV (unpaired t-test). c. In red, same as Figure 2d. In turquoise, normalized histogram of diffusion coefficients of tracks from across the whole nucleus in undifferentiated Halo-SMCHD1^Y353C^ mESCs, kept for 24h with doxycycline. d. Bar plots showing the average percentage of bound molecules of SMCHD1 (3 independent replicates each). Error bars represent the STDEV (unpaired t-test). e. Normalized histogram of diffusion coefficients of tracks from Xi (dark green) or a random region (dark grey) in undifferentiated Halo-SMCHD1^E147A^ mESCs, kept for 24h with doxycycline. f. Normalized histogram of diffusion coefficients of analysed tracks from Xi (light green) or a random region (light grey) in Halo-SMCHD1^E147A^ mESC differentiated for 7 days. g. Bar plot showing the average percentage of bound molecules of SMCHD1 (5 or 4 independent replicates for Halo-SMCHD1 and Halo-SMCHD1^E147A^, respectively) mESCs. Error bars represent the STDEV (unpaired t-test). h. Normalized histogram of diffusion coefficients of tracks from the Xi (turquoise) or a random region (dark grey) in undifferentiated Halo-SMCHD1^Y353C^ mESCs, kept for 24h with doxycycline. i. Normalized histogram of diffusion coefficients of tracks from Xi (light blue) or a random region (light grey) in Halo-SMCHD1^Y353C^ mESCs differentiated for 7 days. j. Bar plot showing the average percentage of bound molecules of SMCHD1 (5 independent replicates each). Error bars represent the STDEV (unpaired t-test). k-l. Average fluorescence recovery curves over the Xi domain (k) or a random nucleoplasmic region (l) in Halo-SMCHD1, Halo-SMCHD1^E147A^ and Halo-SMCHD1^Y353C^ mESCs differentiated for 7 days. The shading represents the STDEV. undiff = undifferentiated, diff = differentiated

Looking at the effect of the introduced mutations on SMCHD1 dynamic behaviour across the Xi domain, for SMCHD1^E147A^, we observed little difference between the population distribution profiles of the mutant and the wild-type protein over the Xi in undifferentiated and differentiated cells (Extended Data Figure 6e, f). This may seem counterintuitive given the absence of enrichment of the mutant protein over the Xi (Figure 4b, Extended Data Figure 5e), but it is key to note that the bound population of SMCHD1^E147A^ in random regions is very similar to that seen at the Xi, both in undifferentiated and differentiated cells (Figure 5e-g). For SMCHD1^Y353C^, binding over the Xi and a random region is unchanged compared to wild-type, both in undifferentiated and differentiated cells and, hence, the mutant protein retains enriched binding at the Xi (Figure 5h-j, Extended Data Figure 6g-h).

Complementary FRAP analysis revealed a highly similar recovery behaviour for SMCHD1^E147A^ within the Xi domain and the nucleoplasm, with the Xi immobile fraction being almost the same as the wild-type protein, but the residence being drastically reduced (Figure 5k, l, Extended Data Figure 7a, c-e). Like SMCHD1^E147A^, SMCHD1^Y353C^ presents a loss of the Xi-specific recovery behaviour observed for wild-type SMCHD1. SMCHD1^Y353C^ recovers to almost pre-bleach levels within the Xi domain and the nucleoplasm, and, importantly, has a much faster turnover compared to the wild-type protein (Figure 5k, l, Extended Data Figure 7b, c-e). This striking decrease in residency on chromatin likely underlies the changes in localization of SMCHD1^Y353C^ at PCH and Xi, given that the population distribution profile assessed by SPT is unchanged. Together, the analysis of GHKL ATPase loss-of-function mutants suggests that ATP hydrolysis is not required for chromatin binding of SMCHD1, but rather affects specificity of chromatin binding, facilitating stable interactions that underlie long-term residence and function at target chromatin sites.

### A gain-of-function mutation in the SMCHD1 GHKL ATPase accelerates Xi recruitment and enhances Xi chromosome compaction

We went on to investigate an additional mutation in the SMCHD1 GHKL ATPase domain, G137E, which results in increased ATPase activity *in vitro*^24,30^ . G137E was identified in a BAMS patient and has been suggested to give rise to a gain-of-function phenotype^24^ . We introduced G137E biallelically at the native *SmcHD1* locus in the wild-type cell line, Halo-SMCHD1 as described for E147A and Y353C (Extended Data Figure 8a, b). In contrast to the other two mutations, G137E does not impair SMCHD1 accumulation at PCH regions or on the Xi (Figure 6a, b, Extended Data Figure 8c). In fact, visual inspection of the cells and quantification of the Xi-associated SMCHD1 intensity show that Xi recruitment is accelerated, notably at days 1-2 following Xist induction (Figure 6b, Extended Data Figure 8c), although we do not observe any effect on the rate of gene silencing (Figure 6c, d).

**Figure 6.**
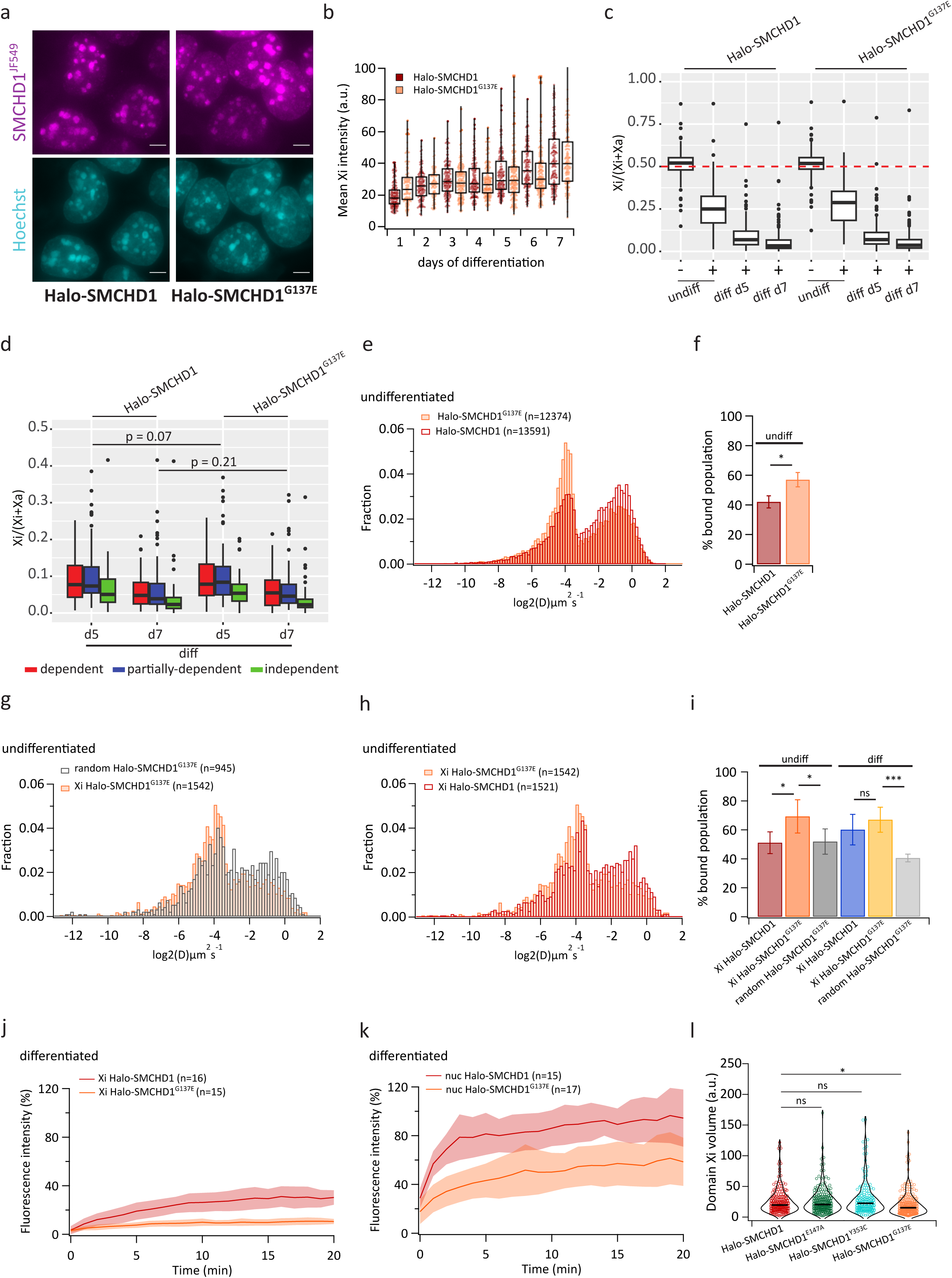
An SMCHD1 GHKL ATPase gain-of-function mutation accelerates Xi association and enhances chromosome compaction. a. Representative live-cell images of indicated mESCs undifferentiated, kept for 24h with doxycycline. Images are maximum-intensity projections, scale bar - 5µm. b. Quantification of SMCHD1 signal underlying CIZ1 domains in indicated cell lines during 7-day differentiation timecourse. Each dot represents a cell. n = 83-200. c. Box plot showing allelic gene expression for X-linked genes in indicated cell lines; uninduced, induced for 24h with doxycycline or differentiated for 5 or 7 days. The boxes represent one or an average of two or three replicates d. Box plot showing allelic X-linked gene expression in indicated cell lines differentiated for 5 or 7 days, with genes split in three categories according to their dependency on SMCHD1 for silencing^13^. The boxes represent an average of two or three replicates. Statistical analysis - paired t test. e. In red, same as Figure 2d. In orange, normalized histogram of diffusion coefficients of tracks from across the whole nucleus in undifferentiated Halo-SMCHD1^G137E^ mESCs, kept for 24h with doxycycline. f. Bar plots showing the average percentage of bound molecules of SMCHD1 (3 independent replicates each). Error bars represent the STDEV (unpaired t-test). g. Normalized histogram of diffusion coefficients of tracks over Xi (orange) or a random region (dark grey) in undifferentiated Halo-SMCHD1^G137E^ mESCs, kept for 24h with doxycycline. h. In red, same as in 3a. In orange, same as in 6g. i. Bar plot showing the average percentage of bound molecules of SMCHD1 (5 or 4 independent replicates for Halo-SMCHD1 and Halo-SMCHD1^G137E^ mESCs, respectively). Error bars represent the STDEV (unpaired t-test). j-k. Average fluorescence recovery curves over the Xi domain (j) or a random nucleoplasmic region (k) in indicated cell lines differentiated for 7 days. The shading represents the STDEV. l. Violin plot showing the volume of the H3K27me3 domain in indicated cell lines differentiated for 7 days. Each dot represents a cell. n = 209 (Halo-SMCHD1), 239 (Halo-SMCHD1^E147A^), 178 (Halo-SMCHD1^Y353C^), 259 (Halo-SMCHD1^G137E^) (unpaired t-test). undiff = undifferentiated, diff = differentiated

SPT analysis of SMCHD1^G137E^ across the nucleus reveals an increased population of bound molecules (Figure 6e, f, Extended Data Figure 8d, e). Over Xi domains, compared to a random region, SMCHD1^G137E^ has a markedly larger bound population in both undifferentiated and differentiated cells (Figure 6g, i, Extended Data Figure 8f). Moreover, the bound population of SMCHD1^G137E^ over Xi in undifferentiated cells is more prominent than that seen in wild-type cells (Figure 6h, i), aligning with the observed tendency of the mutant to accumulate on Xi more rapidly (Figure 6b,). The bound population over the Xi is slightly increased in differentiated cells, but not to a significant level (Figure 6i, Extended Data Figure 8g). In FRAP experiments, SMCHD1^G137E^ recovers more slowly and to a lesser extent compared to wild-type SMCHD1, both in the Xi territory and the nucleoplasm. Importantly, the difference in dynamic behaviour between the two regions observed in the wild-type cells is retained, as determined by FRAP and SPT (Figure 6j, k, Extended Data Figure 9a-d).

SMCHD1 has been reported to promote Xi compaction in human cells^18^, and in a final series of experiments we tested if the different ATPase domain mutations affect Xi volume. To achieve this, we used the Xi-associated histone modification H3K27me3 as a proxy for Xi territory volume, as described previously^44^. No differences in Xi volume were observed for the ATPase loss-of-function mutations SMCHD1^E147A^ and SMCHD1^Y353C^. However, we observed a significant reduction in Xi volume at day 7 of differentiation with SMCHD1^G137E^ (Figure 6l, Extended Data Figure 9e). This result links ATPase activity and Xi compaction, which contextualises the important role for SMCHD1 in XCI.

## Discussion

Our newly developed XX mESC model for live-cell imaging and SPT during establishment of XCI has provided valuable insights into SMCHD1 target site selectivity and its functional interaction with chromatin, including the demonstration that SMCHD1 associates with PCH domains in most cells, and the critical role for LRIF1 in chromatin loading, at both PCH and the Xi. The observation that the majority of SMCHD1 molecules are diffusive in the *Lrif1*^-/-^ cells, unlike in wild-type cells where almost half are chromatin-bound, suggests that the nuclear-wide bound population corresponds in large part to SMCHD1 interacting with H3K9me3-marked heterochromatic regions. H3K9me2-marked heterochromatin may also be a target given the HP1 chromodomain also binds to this histone mark^45,46^. Whilst LRIF1 function accounts for the majority of bound SMCHD1 in mESCs, some PCH domains retain SMCHD1 enrichment in a small proportion of *Lrif1^-/-^*cells, indicating an alternative mechanism for PCH localisation. The basis for this effect is not known but could indicate a pathway that functions at a specific cell cycle stage, or a second adaptor protein that is present only in a small subset of cells.

Involvement of LRIF1 in SMCHD1 recruitment to Xi was surprising given prior evidence that LRIF1 is not required to maintain SMCHD1 on Xi in differentiated somatic cells^18,26^, and moreover, evidence demonstrating a critical role for PRC1-mediated H2AK119ub1 and progression of cellular differentiation for Xi recruitment^11,19^ . LRIF1 in contrast functions as an adapter linking SMCHD1 to H3K9me3 modified chromatin^18,26^. Accordingly, we speculate that a baseline level of chromatin bound SMCHD1 recruited via LRIF1-mediated recognition of H3K9me3 at constitutive heterochromatic regions (PCH and interstitial chromosome arm regions) facilitates stable accumulation of additional SMCHD1 driven by PRC1-mediated H2AK119ub1. This could result from interaction between prebound and newly recruited SMCHD1, or alternatively, reflect an indirect effect resulting from the activity of pre-bound SMCHD1. This interpretation also helps to explain our finding using live-cell imaging that Xi loading of SMCHD1 occurs at the outset of XCI, and as a progressive continuum thereafter, rather than at a specific differentiation stage as previously thought^10,11^.

It has been proposed that SMCHD1 functions similarly to other GHKL ATPases via a molecular clamp mechanism mediated by the ATP binding-hydrolysis-release cycle ^9,30,31^ (Figure 7a). *In vitro* analysis has shown SMCHD1 has a weak ATPase activity which is probably only able to drive conformational changes within the protein itself, as in other GHKL ATPases, rather than power a motor function such as the cohesin-driven loop extrusion process^31,47^. We speculate that such a conformational change may allow the chromatin-bound protein homodimer to simultaneously bind to a different genomic location and, thus, form long-range chromatin interactions (Figure 7b). This conclusion is further underscored by our analysis of GHKL ATPase mutants where effects on stable association with chromatin correlate with loss- or gain-of-function in XCI.

**Figure 7.**
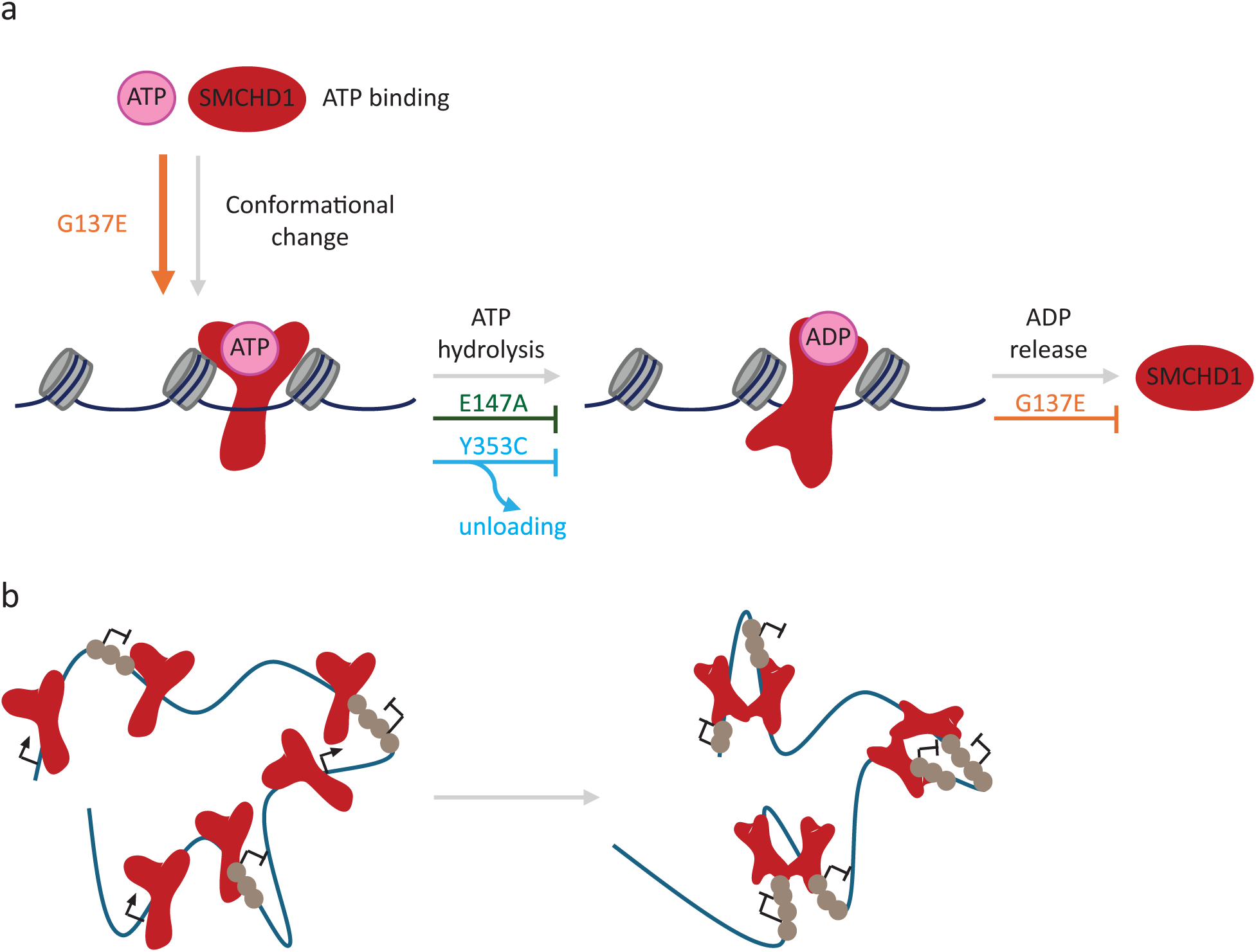
Models for the role of SMCHD1 GHKL ATPase in governing selective chromatin interactions. a. The model invokes that ATP binding to SMCHD1 leads to a conformational change that includes dimerization of the N-terminal GHKL ATPase domains, favours ATP hydrolysis and enables initial loading onto chromatin. Subsequently, ATP hydrolysis takes places, leading to a new protein conformation which favours stable and specific association with chromatin and, potentially, allows distant DNA-bound SMCHD1 molecules to associate with one another (as shown in b.), forming long range chromatin interactions and selective enrichment at specific chromatin regions. Once ADP is released, SMCHD1 unloads from chromatin. The G137E mutation favours ATP binding and, hence, SMCHD1 loading onto chromatin and may also disfavour ADP release and, thus, SMCHD1 dissociation from chromatin. The E147A mutation prevents ATP hydrolysis and locks the protein in the less stably chromatin associated ATP-bound state. The Y353C mutation blocks the catalytic reaction and causes SMCHD1 to unload from chromatin. b. In the context of the Xi, the formation of SMCHD1 mediated long-range chromatin interactions via cycles of ATP binding and hydrolysis may enable spreading of heterochromatic epigenetic marks and silencing factors and, consequently, gene silencing.

In the context of the aforementioned models we interpret the effect of different GHKL ATPase mutations as follows: The E147A mutation, which can bind but not hydrolyse ATP^26,30^, is held in a state that allows engagement with chromatin but prohibits the stable association required for downstream functions, evidenced as reduced residency on chromatin. Consequently, target site selectivity resulting in Xi enrichment of the wild-type protein is lost. The Y353C mutation also abrogates ATPase activity^24,27^ . Our analysis of SMCHD1^Y353C^ indicates a more marked effect than SMCHD1^E147A^ in terms of loss of stable chromatin association. A possible explanation is that the conformational change induced by ATP binding results in the mutant protein disengaging from chromatin rapidly following initial binding, again preventing selective Xi enrichment. Finally, G137E displays increased ADP production and dimerization capacity *in vitro*, suggesting the mutation may facilitate a default conformation which favours ATP binding^24,30^ and, hence, leads to an increase in the bound population of SMCHD1 *in vivo*. Bound SMCHD1^G137E^ has an enhanced residence time on chromatin, potentially due to the G137E-induced conformation disfavouring ADP release and consequently SMCHD1 unloading. Enhanced Xi chromatin association of SMCHD1^G137E^ results in enhanced chromatin compaction. This observation supports that one of the main functions of SMCHD1 is to facilitate long-range chromatin compaction of target loci.

In summary, our findings highlight the key role played by the GHKL ATPase domain for chromatin association and functionality of SMCHD1. In the future, analysis of other mutants coupled to *in vitro* structural and functional investigations of full-length SMCHD1 should provide a more comprehensive understanding. This is key not only to elucidating SMCHD1 function in XCI, but also to understand and develop therapeutic strategies for BAMS and FSHD2 patients.

## Acknowledgements

This work was funded by grants awarded to NB from the Wellcome Trust (215513/Z/19/Z) and UKRI (EP/Y029062/1). SU: Wellcome Trust & Royal Society Sir Henry Dale Fellowship 206159/Z/17/Z and Research Prize Fellowship of the Lister Institute of Preventative Medicine. AS was supported by the Wellcome Trust (209400/Z/17/Z) and the European Research Council (681440) grants awarded to Robert J. Klose. FC received a Clarendon Fund, New College and Department of Biochemistry Scholarship from the University of Oxford. We thank the Klose lab for the Halo-3xNLS, H2B-Halo and Gt(ROSA)26Sor-sgRNA plasmids. We would like to thank staff in Department of Biochemistry and the Dunn School of Pathology imaging facilities for advice and support and members of the Brockdorff lab for critical discussions.

## Data availability

High-throughput raw sequencing data (ChrRNA-seq) as well as key processed data generated in this study are deposited to the National Center for Biotechnology Information’s Gene Expression Omnibus (accession number GSEXXXXXX).

## Materials and methods

### Cell lines

All mouse embryonic stem cells (mESCs) were grown at 37°C, 5% CO_2_ in ES medium: Dulbecco’s Modified Eagle Medium (DMEM; Life Technologies) supplemented with 10% Fetal Calf Serum (ThermoFisher), 0.1mM non-essential amino acids, 2mM L-glutamine, 50μM β-mercaptoethanol, 100 U/ml Penicillin/100 μg/ml Streptomycin (all from Life Technologies) and 1000U/mL Leukemia Inhibitory Factor (LIF) (made in-house). The cells were grown on gelatinized plates, in feeder-dependent conditions. Feeders were generated from mitomycin-inactivated (Sigma Aldrich) SNLPs (STO mouse fibroblasts expressing Neomycin, Puromycin resistance and Lif genes). The main parental cell line for all subsequently derived cell lines is the interspecific (129/sv - Cast/Ei) iXist-ChrX XX mESC line previously derived and described in^33^. Briefly, these cells include a tetracyline-inducible promoter (TetON) on the *M.m. domesticus* (129/sv) Xist allele and a reverse tetracycline-controlled transactivator (rtTA) expressed from the *Tigre* locus. For Xist induction, cells were treated with 1μg/ml doxycycline (Sigma).

### Cloning of plasmids for CRISPR-Cas9 targeting

For the homology vectors (HVs), homology arms of approximately 500-600bp flanking the insertion site were amplified by PCR from mESC genomic DNA using Fast-Start High Fidelity enzyme (Merk Life Science). The Halo-, Snap- and eGFP-tags were amplified from previously generated vectors containing these sequences. The fragments were then ligated together with the backbone plasmid by Gibson Assembly ligation using Gibson Assembly Master Mix (NEB) according to the manufacturer’s guidance. For most HVs, the primers for Gibson Assembly were designed so that they introduced mutations to the PAM recognition sequence of the corresponding sgRNAs and/or mutations to create restriction enzyme (RE) sites required for screening by PCR followed by RE digestion. When introduction of such mutations was not possible through the Gibson Assembly primers, the plasmids obtained were subsequently mutagenized using QuikChange Lightning Site-Directed Mutagenesis kit (Agilent).

Selection of the sgRNAs required for recognition and subsequent double-strand break generation at the target loci by CRISPR-Cas9 was done using the online tool CRISPOR ^48^. Annealed dsDNA oligos matching the chosen sgRNA were cloned into the PX459 V2.0 backbone (Addgene plasmid #62988) following the single-step digestion-ligation Zhang lab protocol^49^. The sgRNA sequences used are listed in Table S1.

The ligated products were transformed into XL10-Gold ultracompetent bacteria (Agilent) and DNA was isolated from single bacterial colonies using the Miniprep kit (Qiagen). The plasmid sequences were checked via Sanger sequencing (SS) (Source BioScience).

**Table S1:**
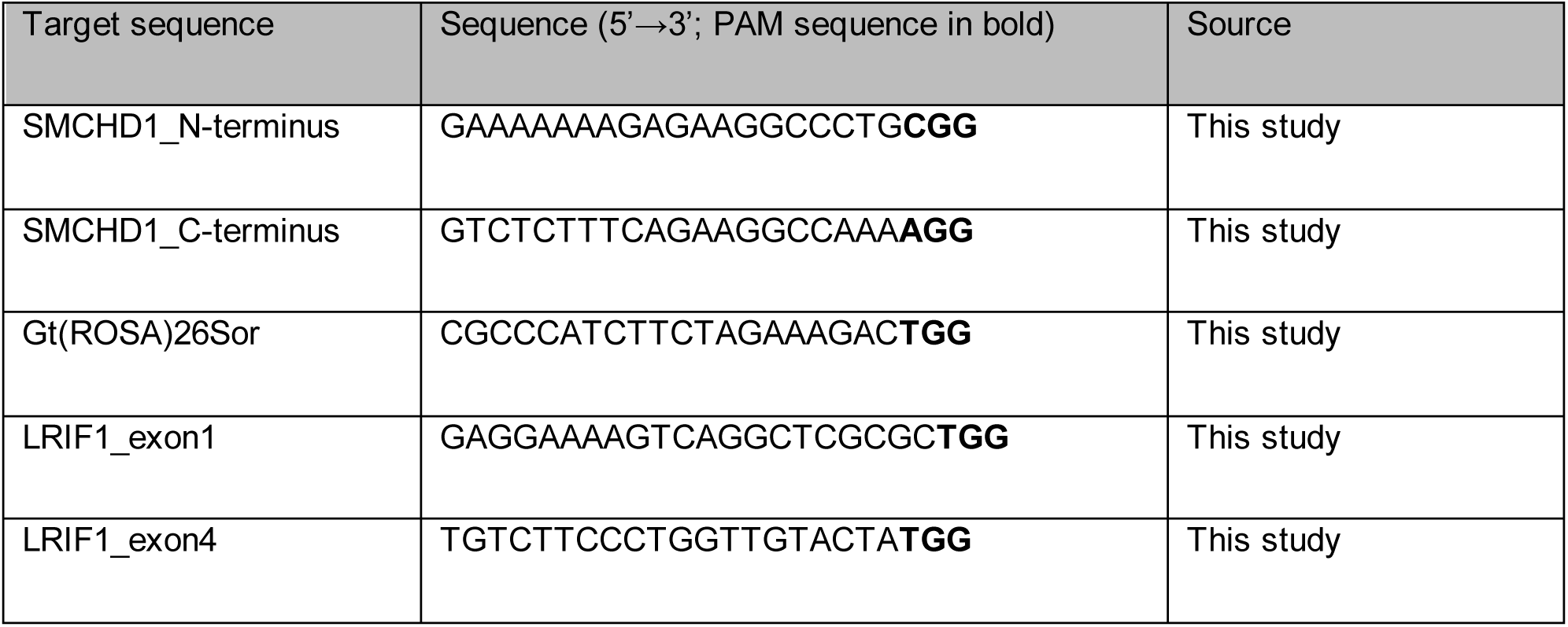
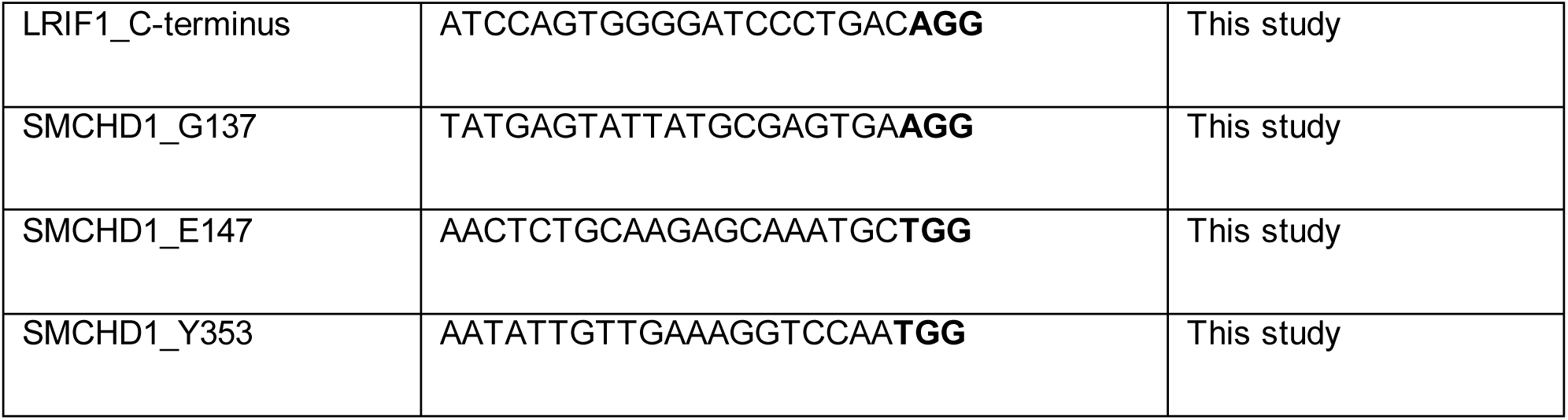
Sequences of sgRNAs used for CRISPR-Cas9 targeting.

### Generation of stable cell lines using CRISPR-Cas9-mediated homologous recombination

A day before transfection, 250 000 cells/well were seeded in 6-well plates on feeders. Cells were transfected using Lipofectamine 2000 (ThermoFisher) according to the manufacturer’s protocol. The HVs were co-transfected with the Cas9-sgRNA plasmids at a molar ratio of 6:1 (2.4µg HV and ∼1µg Cas9-sgRNA plasmid). For the LRIF1 KO cell line, two Cas9-sgRNA plasmids targeting exons 1 and 4 were transfected, each at 1µg. Transfected cells were passaged 24h post-transfection onto gelatinized 90mm Petri dishes with feeders and, 48h post-transfection, puromycin (3-5µg/ml) selection was applied. Cells were kept under selection for 48h followed by exchanging for fresh mESC media every day for a further 7-9 days until individual colonies were picked. The clones were then expanded in 96-well plates for screening. A summary of the cell lines generated is shown in Table S2 and a list of the primers used for screening is shown in Table S3.

**Table S2:** Summary of XX mESC lines derived in this study and associated screening strategies.

| Cell line | Screening strategy |
| --- | --- |
| Halo-SMCHD1 | PCR, SS, WB and live imaging |
| SMCHD1-Snap | PCR, SS, WB and live imaging |
| LRIF1 KO | PCR, SS and WB |
| LRIF1-Halo | PCR, SS, WB and live imaging |
| Halo-SMCHD1<sup>G137E</sup> | PCR followed by RE digestion, SS |
| Halo-SMCHD1<sup>E147A</sup> | PCR followed by RE digestion, SS |
| Halo-SMCHD1<sup>Y353C</sup> | PCR followed by RE digestion, SS |
| H2B-Halo | Live imaging |
| Halo-3xNLS | Live imaging |

**Table S3:**
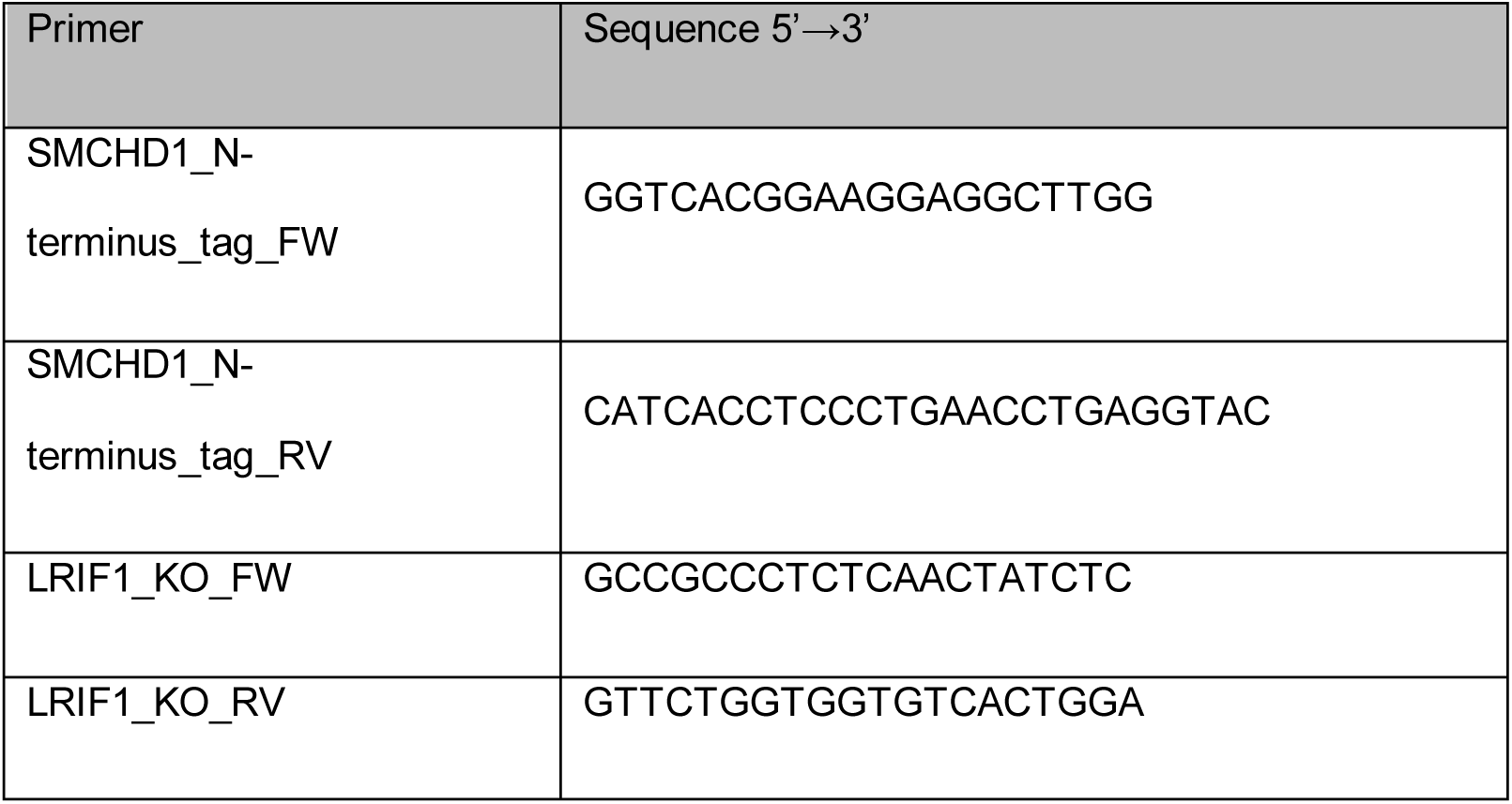
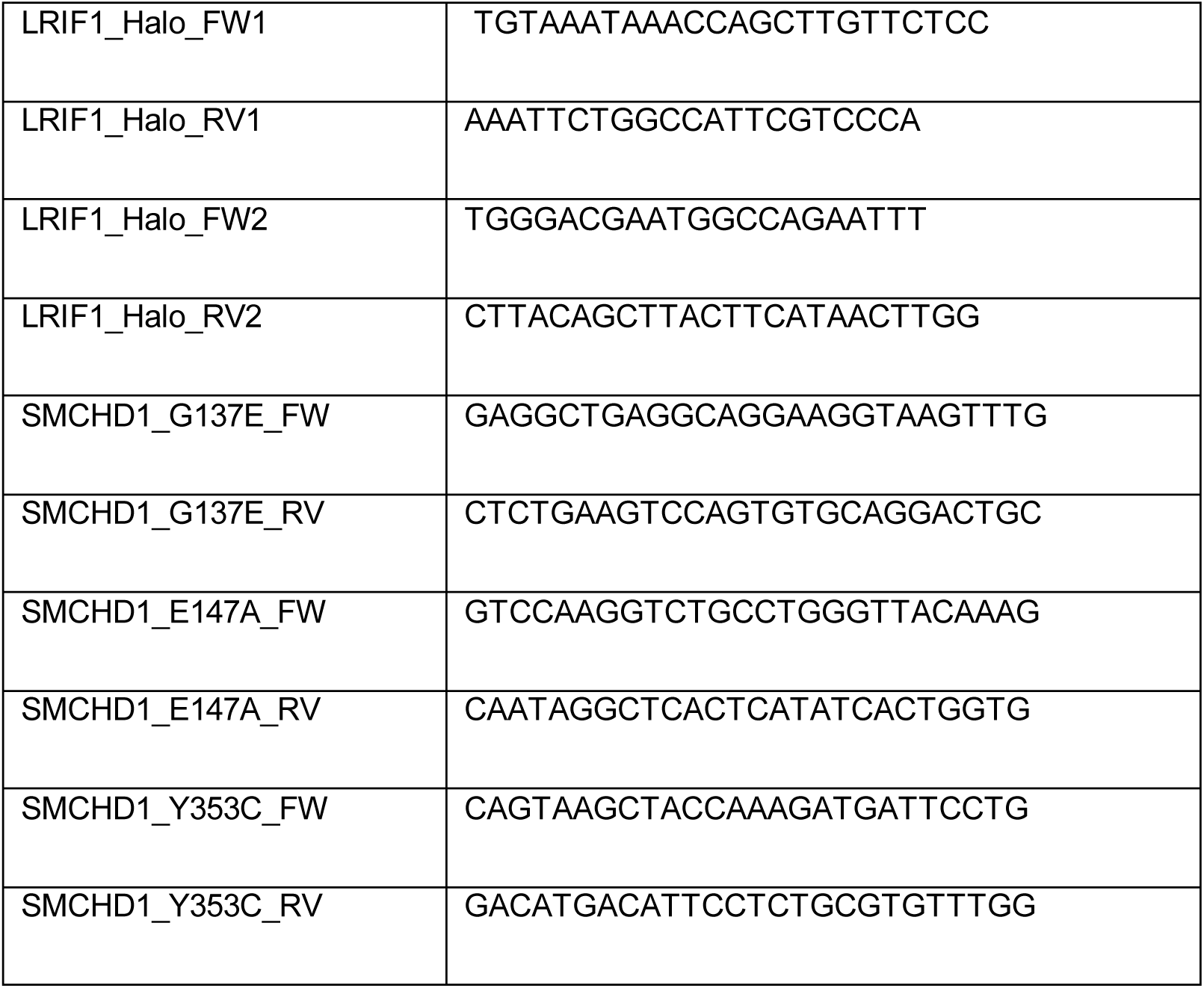
List of primers used for screening of colonies from 96well plate.

### Monolayer (ML) differentiation

For *in vitro* differentiation, the cells were pre-plated for 30 - 40min in non-gelatinized plates to remove feeder cells, due to feeders faster attachment relative to mESCs. The collected cells in suspension, which are primarily mESCs, were counted and plated at a density of 2000 cells/cm^2^ in gelatinised plates. The differentiating cells were grown for up to 7 days. From the pre-plating step onwards, cells were kept in EC10 + dox medium: Dulbecco’s Modified Eagle Medium (DMEM; Life Technologies) supplemented with 10% Fetal Calf Serum (ThermoFisher), 0.1mM non-essential amino acids, 2mM L-glutamine, 50μM β-mercaptoethanol, 100 U/ml Penicillin/100 μg/ml Streptomycin (all from Life Technologies) and 1μg/ml doxycycline (Sigma).

### Single particle tracking (SPT)

Cells were plated on gelatinised 35 mm µ-dishes containing a #1.5 polymer coverslip (Ibidi) two days before imaging in the case of differentiated cells or one day before imaging for undifferentiated cells. In the latter case, after plating, 1μg/ml doxycycline was added to the dish and the cells were imaged 24h later. Halo-PA-JF549 ligand (HHMI Janelia)^50^ was used for labelling at 100nM for 15min at 37 ℃. Cells were then washed twice with PBS (Sigma) and incubated for 30min in Fluorobrite DMEM (ThermoFisher) supplemented with 10% Fetal Calf Serum (ThermoFisher) and 1μg/ml doxycycline. After three more washes with PBS, cells were imaged in Fluorobrite DMEM supplement as above and additionally with 30mM HEPES pH 7.4 to maintain the pH of the sample medium.

SPT experiments were carried out with a custom TIRF/HILO microscope, described previously^51^, with an EMCCD camera (Andor), laser module (iChrome MLE MultiLaser engine, Toptica Photonics), 100x 1.4NA objective (Olympus) and translational module (ASI) which allows for adjustment of the angle of the excitation beam between epi- and HILO-illumination to achieve a high signal-to-noise ratio.

To visualise both diffusing and bound molecules, movies of 4000 frames, with 15ms/frame, were acquired with continuous 561nm laser excitation at 22mW intensity and 405nm laser excitation at varied intensities to maintain a low density of fluorescent particles. Importantly, for the Xi-focused acquisitions, an image of the Xi region(s) was taken before and after the movie acquisition with 488nm laser excitation at 2mW and 20ms camera exposure. For whole nucleus datasets, at least 20 movies were taken, each containing about three cells, and at least three replicates were performed for each timepoint studied. For Xi-focused datasets, 50 movies were taken, each containing at least one Xi domain, and at least four replicates were performed for each timepoint studied. The precise total number of movies/Xi/random regions analysed for each timepoint is shown in Table S4.

**Table S4:** Summary of no. of movies/regions analysed for each cell line using SPT.

| Cell line | Timepoint | No. of analysed movies/regions |
| --- | --- | --- |
| Halo-SMCHD1 | Undifferentiated (24h doxycycline) | 60 movies, 343 Xi/random regions |
|  | Differentiated (7 days) | 64 movies, 307 Xi/random regions |
| LRIF1 KO | Undifferentiated (24h doxycycline) | 64 movies, 272 Xi/random regions |
|  | Differentiated (7 days) | 66 movies, 235 Xi/random regions |
| Halo-SMCHD1<sup>E147A</sup> | Undifferentiated (24h doxycycline) | 64 movies, 262 Xi/random regions |
|  | Differentiated (7 days) | 66 movies, 242 Xi/random regions |
| Halo-SMCHD1<sup>Y353C</sup> | Undifferentiated (24h doxycycline) | 66 movies, 246 Xi/random regions |
|  | Differentiated (7 days) | 66 movies, 195 Xi/random regions |
| Halo-SMCHD1<sup>G137E</sup> | Undifferentiated (24h doxycycline) | 64 movies, 262 Xi/random regions |
|  | Differentiated (7 days) | 66 movies, 242 Xi/random regions |
| H2B-Halo | Undifferentiated (24h doxycycline) | 22 movies |
| Halo-3xNLS | Undifferentiated (24h doxycycline) | 20 movies |

### SPT data analysis

Single particle tracking datasets were analysed predominantly in MATLAB (MathWorks) using Stormtracker software^52^, unless otherwise specified. Localization of single particles in each frame of the acquired movies was achieved with a precision of 25 nm by fitting an elliptical Gaussian point spread function to the detected molecules. The detection of the molecules was done based on an intensity threshold fixed for all movies taken within the same acquisition session but adjusted as needed from session to session after manually checking the detection efficiency. Once localized, the molecules found within a radius of 768 nm in consecutive frames of the movie were linked to form tracks; single frame gaps were allowed, to take into account for spontaneous fluorophore blinking. The tracks which contained a minimum of four frames were then used to calculate their apparent diffusion coefficient D*.

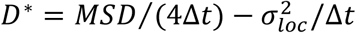

where Δt = 15.26 msec, σ_loc_ = 40 nm and MSD is the mean-squared displacement:

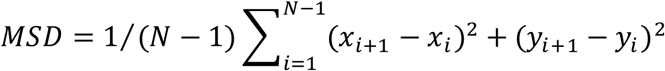

where N ≥ 5 as only tracks with at least four frames are considered.

The log2 values corresponding to all the tracks found across all movies and all replicates for each timepoint analysed were displayed as a histogram using Igor Pro 9 (WaveMetrics); the histograms were normalised to the number of tracks analysed in each timepoint.

Tracks with a log2(D*) smaller than -3 were considered to represent bound molecules and the bound population was calculated as a percentage of bound tracks relative to the total analysed tracks for that timepoint. For Extended Data Figure 2g, Gaussian fitting and SPOT-ON kinetic modelling were done as previously described^42,43^.

For the Xi-focused datasets, a filtering step was added to attribute the localized molecules found within the Xi territory prior to linking these localized molecules into tracks. To do this, firstly, the two images of the Xi region, taken before and after the movie acquisition, were merged using ImageJ/FiJi. If the region was found to have significantly moved during the acquisition, that particular field of view was discarded from further analysis. A custom ImageJ/FiJi script was used to manually select the Xi region in each of the merged images, as well as a random region of the same size as the Xi region but drawn somewhere else in the nucleus. The script outputs files containing the xy coordinates of these regions. These files were inputted in a second custom-made R script used to filter the localized single molecules, keeping only the molecules present in each of the selected regions. The filtered molecules were then merged into tracks which were analysed as described above.

### Live cell imaging

Cells were plated in gelatinised µ-slide 8 wells or 35 mm µ-dishes with a polymer coverslip (Ibidi) at least four hours before imaging. Before imaging, cells were labelled with Halo-JF549 ligand (Promega) at 100nM for 30min at 37 ℃. Cells were then incubated for 30min in growing media and subsequently washed twice with PBS (Sigma). Cells were imaged in Fluorobrite DMEM (ThermoFisher) supplemented with 10% Fetal Calf Serum (ThermoFisher) and, if needed, 1μg/ml doxycycline and/or 0.1µg/ml Hoechst (ThermoFisher). Live cell imaging was performed using an Olympus IX83 microscope equipped with a 63x 1.4NA objective, a 1200 x 1200 px^2^ sCMOS camera (Photometrics 95B) and a humidified chamber with carbon dioxide atmosphere at 37℃. The microscope was operated via CellSens software. When needed, an additional magnifying 1.6x or 2x lens was used in front of the camera leading to a final pixel size of 114.4 nm or 91.5nm, respectively. z-stacks were captured every 0.5 µm, using 10-250ms exposure at 3-50% laser power depending on the channel. All acquired images were analysed and processed using ImageJ/Fiji.

### Time course setup and analysis

For live imaging of the 7-day ML differentiation timecourse, all analysed cells were seeded for differentiation at the same time. Each day of the timecourse, cells were plated on 8 well µ-slides with a polymer coverslip (Ibidi) four hours before imaging. The images were acquired with the Olympus IX83 microscope using the 2x lens in front of the camera. The analysis was performed in ImageJ/FiJi using maximum-intensity projections of the acquired images. A binary mask of the Xi domains was created by thresholding the green channel images representing eGFP-CIZ1 signal. Each red channel image representing Halo-JF549-SMCHD1 signal was multiplied by its corresponding binary mask and then the Analyse Particles function was used to determine the mean SMCHD1 signal intensity in each analysed Xi domain. The values for each day were plotted as a box plot using Igor Pro 9.

### Fluorescence recovery after photobleaching (FRAP)

Differentiating cells were plated on gelatinised 35 mm µ-dishes with a polymer coverslip (Ibidi) two days before imaging. Trolox 1mM (Sigma-Aldrich) was added to the media 24h before imaging to reduce phototoxicity as a result of prolonged imaging. Before imaging, cells were labelled as described above with Halo-JF549 ligand and imaged in Fluorobrite DMEM supplemented with 10% Fetal Calf Serum and 1µg/ml doxycycline. Live-cell timecourse imaging was performed using an Olympus SoRa spinning disc confocal equipped with a 63x 1.4NA objective, sCMOS camera (Photometrics), 405nm FRAP module and a humidified chamber with carbon dioxide atmosphere at 37℃. The microscope was operated using the CellSens software. The ROIs, either Xi or nucleoplasmic regions, were manually selected using the two-point circle measuring tool and photobleached with one bleach pulse of 40% laser intensity, following acquisition of two pre-bleached images with a delay of 1 minute between them. After photobleaching, images were taken every minute for a total of 20 minutes. Each image was taken with 100ms camera exposure with 10% 568nm laser intensity and consisted of a z-stack of 33 slices captured every 0.5µm. The acquisition was performed on three different days for each cell line analysed.

### FRAP analysis

Maximum intensity z-projections were acquired and the fluorescence loss due to photobleaching across the timecourse was corrected for using the bleach correction plugin in ImageJ/FiJi, fitting to an exponential decay. The fluorescence intensity measured within the bleached region was normalised to the background intensity, measured outside of the cell, and the fluorescence recovery was then represented as a percentage relative to the fluorescence intensity pre-bleaching. Curves showing the averaged recovery of all cells analysed for each condition as a function of time were plotted using Igor Pro 9. The curves were then fitted with a single exponential function in Igor Pro 9 in order to determine the immobile population and residency time.

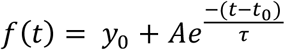

### Immunofluorescence

Cells were plated on gelatinised 35 mm µ-dishes two days before fixation. After two washes with PBS, cells were fixed in 2% paraformaldehyde for 15min at RT and permeabilized in 0.4% Triton X-100 for 5min at RT. Cells were briefly washed with PBS, blocked for three periods of 2min with 0.2% fish gelatine in PBS and incubated with primary antibody (anti-SMCHD1 1:1000 (made in-house), anti-H3K27me3 1:1000 (Active Motif, # 61017) diluted in fish gelatine solution with 5% normal goat serum for 2h at RT in a humid chamber. Samples were washed three times with fish gelatine solution and incubated for 1h at RT with secondary antibody (Alexa Fluor 568 goat anti-mouse IgG, Alexa Fluor 488 goat anti-rabbit IgG) 1:500 diluted in fish gelatine solution. Following two washes with fish gelatine solution and one wash with PBS, Vectashield with DAPI was added and the µ-dish lid was locked with parafilm. Cells were analysed on the Olympus IX83 microscope.

### H3K27me3 domain size analysis

Z-stack images taken with the Olympus IX83 microscope were analysed using the 3D Objects Counter function in Fiji/ImageJ. For each image, a threshold was set in order to highlight the H3K27me3 focus in each cell and the volume of each focus was chosen as an output. The object map was cross-checked against the original image to make sure the foci have been correctly identified. For each cell line, the volume values were plotted as a violin plot using Igor Pro 9.

### Chromatin-associated RNA extraction and sequencing (ChrRNA-seq)

mESCs growing without feeders or ML-differentiated cells were pelleted from confluent 90mm dishes, washed once with PBS and snap-frozen for storage at -80℃. The pellets were resuspended in HLBN buffer (0.01M NaCl, 2.5mM MgCl_2_, 10mM Tris pH 7.5, 0.1% NP-40). Then the cells were underlaid with HLBN buffer supplemented with 24% sucrose and nuclei were isolated through centrifugation (1000g 5min at 4℃). Nuclei were resuspended in NUN1 (75mM NaCl, 20mM Tris pH 7.9, 0.5mM EDTA, 50% glycerol, 0.1mM DTT) and then lysed with NUN2 (0.3M NaCl, 20mM HEPES pH 7.6, 7.5mM MgCl_2_, 0.2mM EDTA, 1M urea, 1% NP-40, 0.1mM DTT) by incubating 15min on ice and occasionally vortexing. The chromatin was isolated by centrifugation (15000rpm 10min at 4℃) and resuspended in HSB buffer (0.5M NaCl, 10mM Tris pH 7.5, 10mM MgCl_2_). The samples were DNAse (LifeTech) treated at 37℃, shaking (1400rpm) until the chromatin pellet dissolved. Samples were then treated with 0.2% SDS and 0.4mg/ml proteinase K for 30min at 37℃, shaking (1400rpm). To obtain the chromatin-associated RNA, two rounds of standard TRIzol- (Invitrogen) chloroform extraction with overnight isopropanol precipitation were performed. Between the two rounds, a second DNAse treatment was performed for 30min at 37℃, shaking (1400rpm). After the second round, the RNA pellets were resuspended in RNase-free water and their concentration was measured by NanoDrop (ThermoFisher). 100-500ng of RNA was used for library preparation with the Illumina TruSeq stranded total RNA kit (RS-122-2301). The pooled libraries were sequenced by 2 x 81 paired-end sequencing using Illumina NextSeq500 (FC-404-2002).

### ChrRNA-seq analysis

The analysis of the ChrRNA-seq datasets was done as previously described ^33^. Briefly, raw fastq files were mapped using bowtie2^53^ (v2.3.2) to rRNA and the rRNA-mapped reads were discarded. Subsequently, the unmapped reads were aligned to an N-masked mm10 genome with STAR^54^ (v2.4.2a or v2.7.9a) with parameters “*-- twopassMode Basic --outSAMstrandField intronMotif --outSAMattributes All -- outFilterMultimapNmax 1 --outFilterMismatchNoverReadLmax 0.06 --alignEndsType EndToEnd*”. Unique alignments were then split into the two alleles, Cast and 129S, using SNPsplit^55^ (v0.4.0dev). featureCounts program^56^, with parameters “*-t transcript - g gene_id -s 2*”) was used to count the allelic and unsplit reads that overlap with genes. Samtools^57^ (v1.16.1) was used to filter and sort the alignments. Only genes with at least 10 SNP-covering reads across samples for comparisons were used to calculate the allelic ratio Xi/(Xi+Xa) (Xi represents the inactive allele which here is the 129 one, while Xa represents the active allele which here is the Cast one). The genes were split into three categories according to their dependency on SMCHD1 for silencing as previously defined in^13^. Boxplots were generated using R package ggplot2 and p-values were calculated using Paired *t* test. The designed mutations (i.e. G137E, E147A, Y353C) for SMCHD1 are confirmed by the ChrRNA-seq reads and are visualised using IGV.

### Whole cell lysates, RIPA extracts, nuclear extracts and western blot

Whole cell lysates were obtained from mESCs growing without feeders, by washing the cells with PBS and then lysing them with SMASH buffer (0.125M Tris pH 6.8, 11.6% glycerol, 4% SDS, 0.002% bromophenol blue, 10% β-mercaptoethanol). The lysates were snap-frozen and stored at -80℃ until use.

For nuclear extracts, mESCs growing without feeders were pelleted, washed with PBS, snap-frozen on dry ice and stored at -80℃. The pellets were resuspended in hypotonic buffer A (10mM HEPES pH 7.9, 1.5mM MgCl_2_, 10mM KCl) supplemented with 0.5mM DTT, 0.5mM PMSF and 1x complete protease inhibitors (Roche) and incubated on ice for 10min. Cells recovered by centrifugation, were lysed in buffer A with 0.1% NP-40 supplemented with 5mM DTT, 0.5mM PMSF and 1x complete protease inhibitors and incubated on ice for 10min. The nuclei were then isolated by centrifugation, washed once with PBS supplemented with 1x complete protease inhibitors, and resuspended in high salt buffer C (5mM HEPES pH 7.9, 26% glycerol, 1.5mM MgCl_2_, 0.2mM EDTA, 350mM NaCl) supplemented with 0.5mM DTT and 1x complete protease inhibitors. The extraction was done on ice for 1h and the nuclei were then pelleted with the supernatant kept as the nuclear extract which was snap-frozen and stored at -80℃ until use. The concentration of the extract was determined using the Bio-Rad Bradford assay according to the manufacturer’s protocol. To analyse the chromatin insoluble fraction, the last nuclear pellet obtained was resuspended in SMASH buffer, snap-frozen and stored at -80℃ until use.

The extracts obtained through one of the methods above were separated on a home-made SDS-PAGE separating gel with a stacking gel or a commercial NuPAGE gel (ThermoFisher). The proteins were transferred onto a nitrocellulose membrane (Bio-Rad) using the Bio-Rad Trans-Blot Turbo Transfer system, with High MW or Mixed MW preprogrammed protocols depending on the protein size. Membranes were blocked for 1h at RT in PBST (PBS with 0.1% Tween 20) with 5% milk and incubated with primary antibodies (summarised in Table S5) diluted in blocking solution, overnight shaking at 4°C. Following washes with PBST, membranes were incubated for 1h at RT with secondary antibodies conjugated to horseradish peroxidase (Amersham) diluted 1:2000 in blocking solution. Membranes were washed with PBST and the bands were revealed using enhanced chemiluminescence (ECL) solution (ThermoFisher) and developed on chemiluminescent sensitive photographic film (Amersham).

**Table S5:**
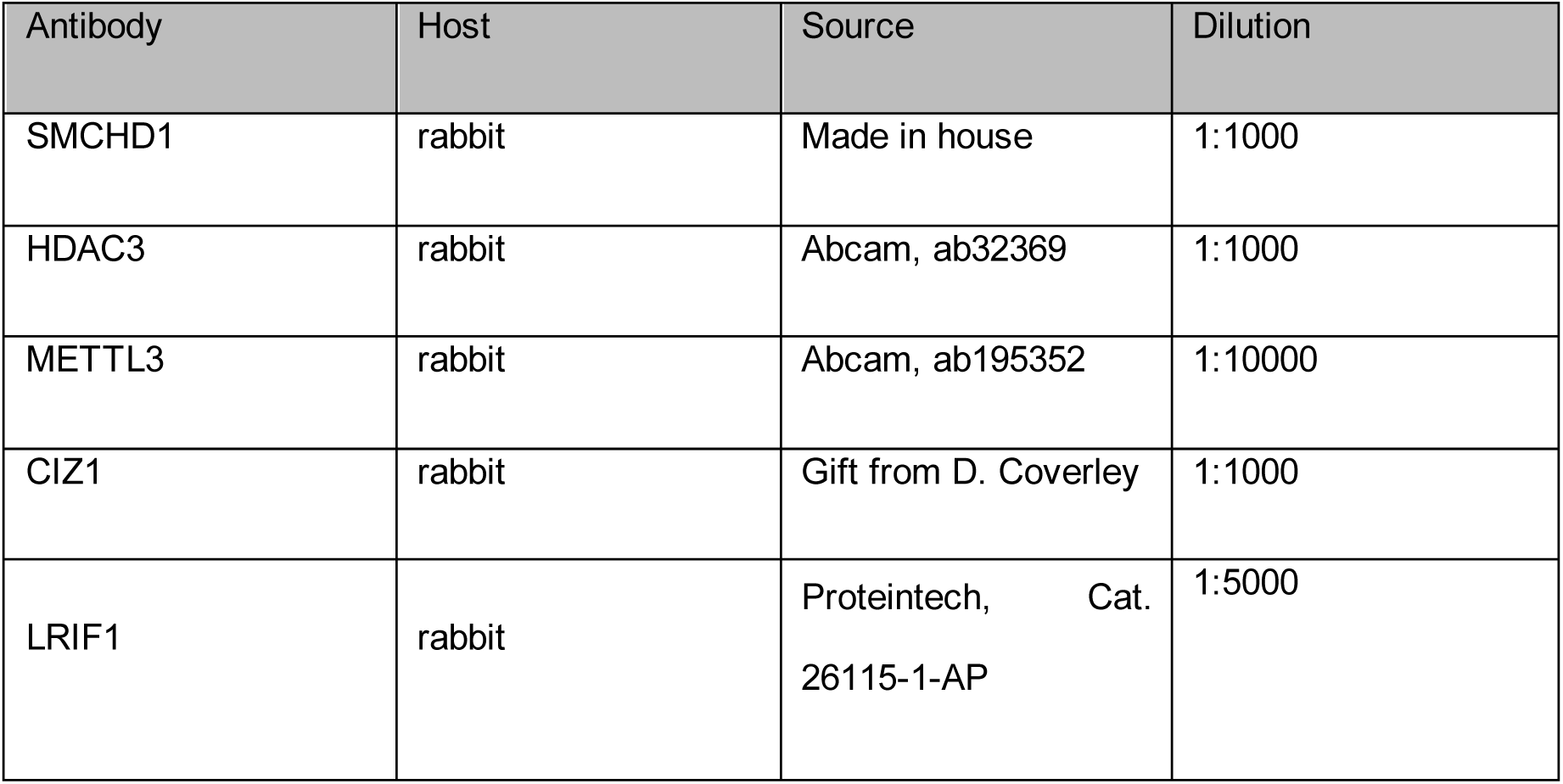
Antibodies used for Western Blot.

| Antibody | Host | Source | Dilution |
| --- | --- | --- | --- |
| SMCHD1 | rabbit | Made in house | 1:1000 |
| HDAC3 | rabbit | Abcam, ab32369 | 1:1000 |
| METTL3 | rabbit | Abcam, ab195352 | 1:10000 |
| CIZ1 | rabbit | Gift from D. Coverley | 1:1000 |
| LRIF1 | rabbit | Proteintech, Cat. 26115-1-AP | 1:5000 |

---

**Extended Data Figure 1.**
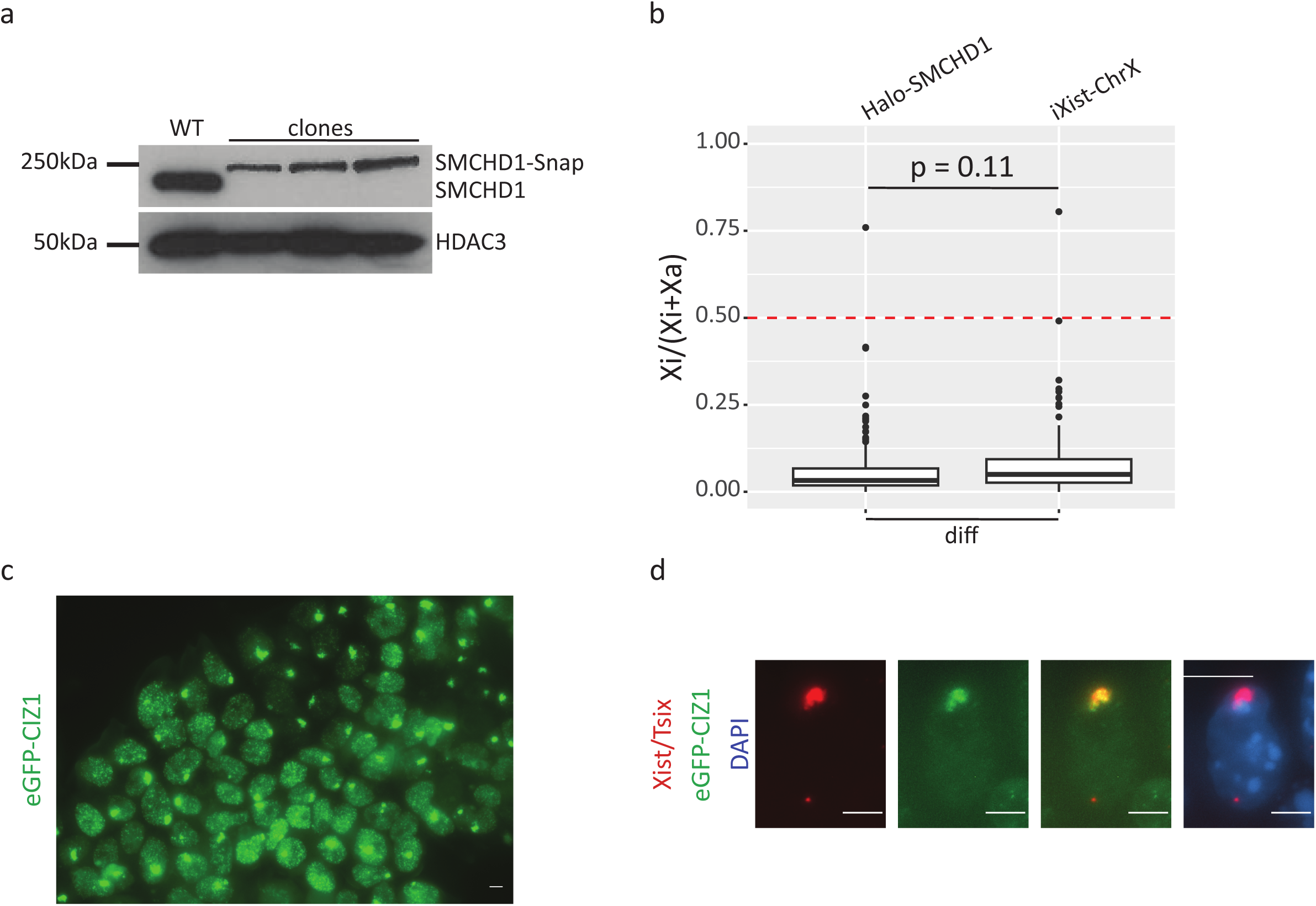
Validation of a model system used to study the dynamic behaviour of SMCHD1. a. Western Blot showing the level of expression of SMCHD1-Snap in three independent clones. b. Representative image of the proportion of Halo-SMCHD1 mESCs showing CIZ1-marked Xi domain following 24h doxycycline treatment. Image is a maximum-intensity projection, scale bar - 5µm. c. Representative images of Xist FISH and eGFP-CIZ1 signal in fixed samples of Halo-SMCHD1 mESC line following 24h doxycycline addition. Images are maximum-intensity projections, scale bar - 5µm. d. Box plot showing allelic ChrRNA-seq analysis of X-linked gene silencing in Halo-SMCHD1 clone G2 and iXist-ChrX mESCs. Boxes are averaged from two or three replicate differentiation experiments. Statistical analysis - paired t-test.

**Extended Data Figure 2.**
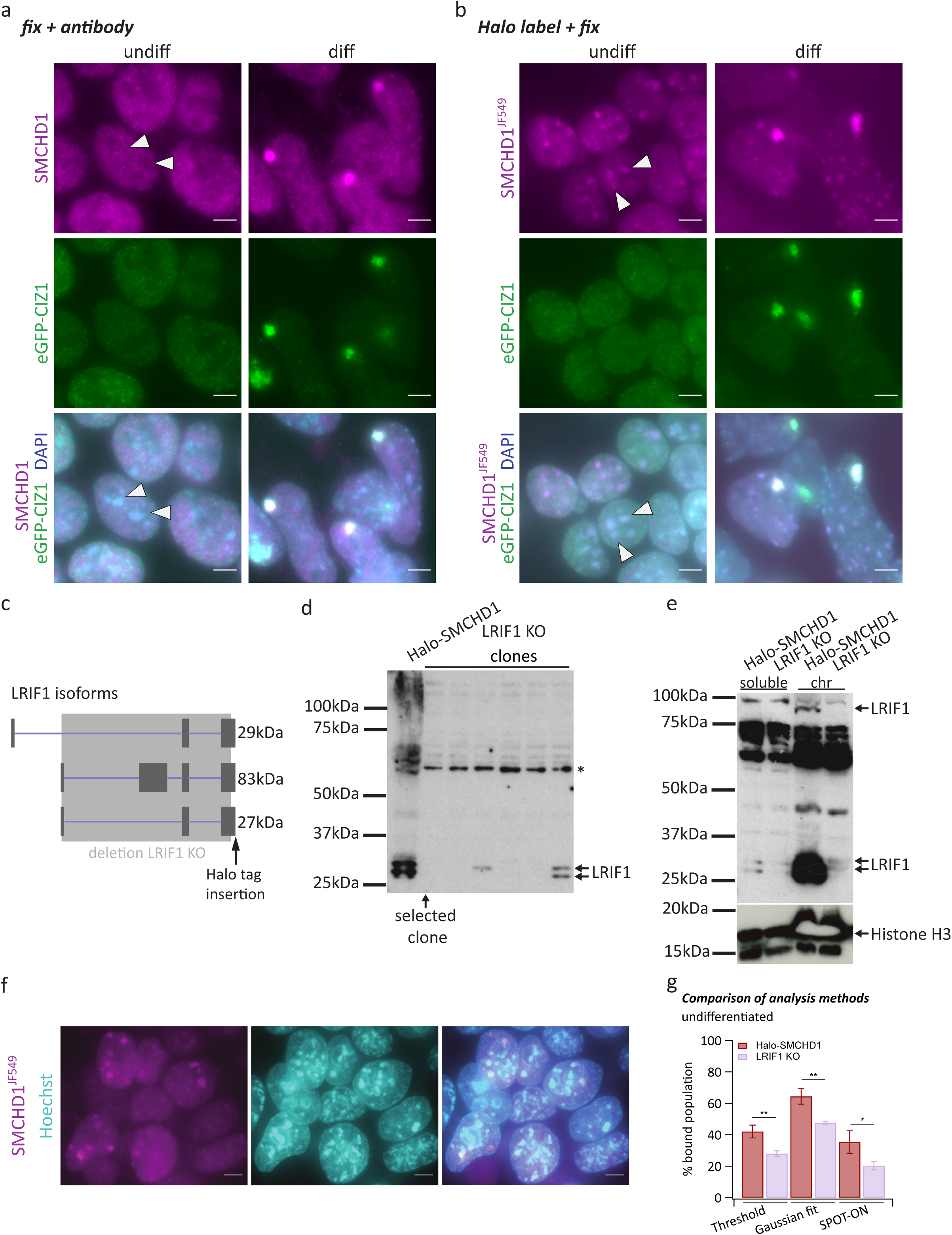
Characteristics of SMCHD1 enrichment over PCH domains. a. Representative images of SMCHD1 immunostaining in fixed samples from Halo-SMCHD1 mESCs undifferentiated or differentiated for 7 days. Arrowheads point towards examples of PCH domains. b. Representative images of Halo labelling prior to sample fixation from Halo-SMCHD1 mESCs undifferentiated or differentiated for 7 days. Arrowheads point towards examples of PCH domains. c. Schematic showing the three *LRIF1* isoforms and the engineering strategies for KO and C-terminal tagging. d. Western Blot showing the expression of LRIF1 in Halo-SMCHD1 mESC (first lane) and from potential Lrif1^-/-^ clones obtained (subsequent 6 lanes). * indicates non-specific band. e. Western Blot from nuclear extract showing the soluble fraction in the first two lanes and the insoluble chromatin fraction in the last two lanes. Note that the amount of protein in the chromatin fraction samples was not quantified but non-specific bands on the blot suggest there is roughly the same amount loaded between the Halo-SMCHD1 and LRIF1 KO samples. f. Representative live-cell image of Lrif1^- /-^ mESCs showing retention of SMCHD1 at some PCH regions. All images are maximum-intensity projections, scale bar - 5µm. g. Bar plots showing the average percentage of bound molecules of SMCHD1 the indicated mESC lines (3 independent replicates each), using three different analysis methods. Error bars represent the STDEV (unpaired t-test).

**Extended Data Figure 3.**
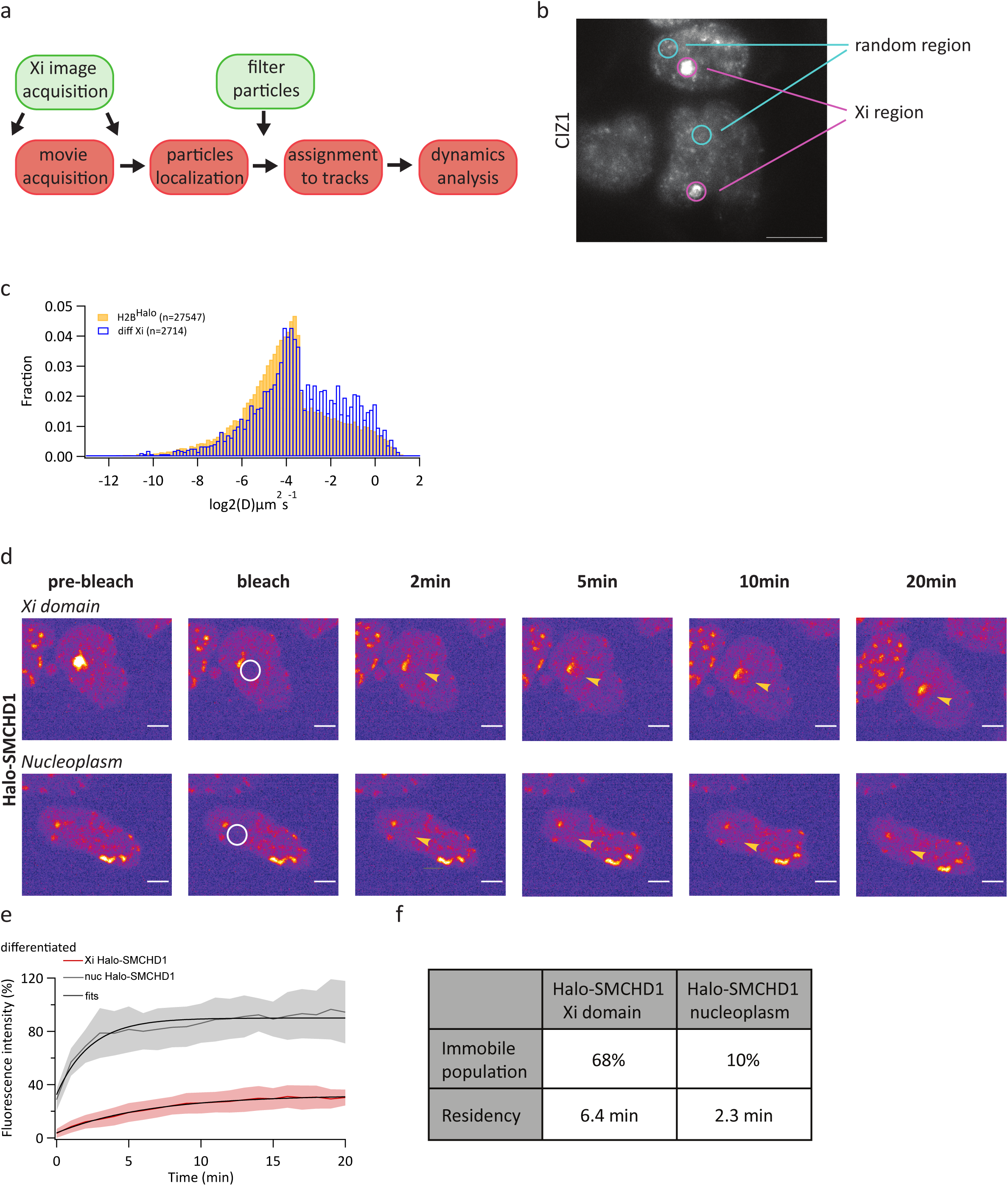
Characterisation of SMCHD1 Xi-specific dynamic behaviour. a. Schematic of Xi-focused SPT analysis. b. Representative image of Xi domains marked by CIZ1 accumulation, taken on the TIRF/HILO microscope used for SPT. Two examples of areas used for control random regions are also shown. Scale bar - 10µm. c. In blue, same as in Figure 3b. In yellow, normalized histogram of diffusion coefficients of tracks from across the whole nucleus in histone H2B-Halo mESCs. d. Representative images from a FRAP experiment using Halo-SMCHD1 mESC differentiated for 7 days. The bleached region is marked by a white circle on the bleach snapshot and by a yellow arrow in the post-bleach snapshots. Scale bar - 5µm. e. Single exponential fits of the average fluorescence recovery curves over the Xi domain or a random nucleoplasmic region in Halo-SMCHD1 mESC. f. Summary of the immobile population and residency values extracted from the fits shown in e.

**Extended Data Figure 4.**
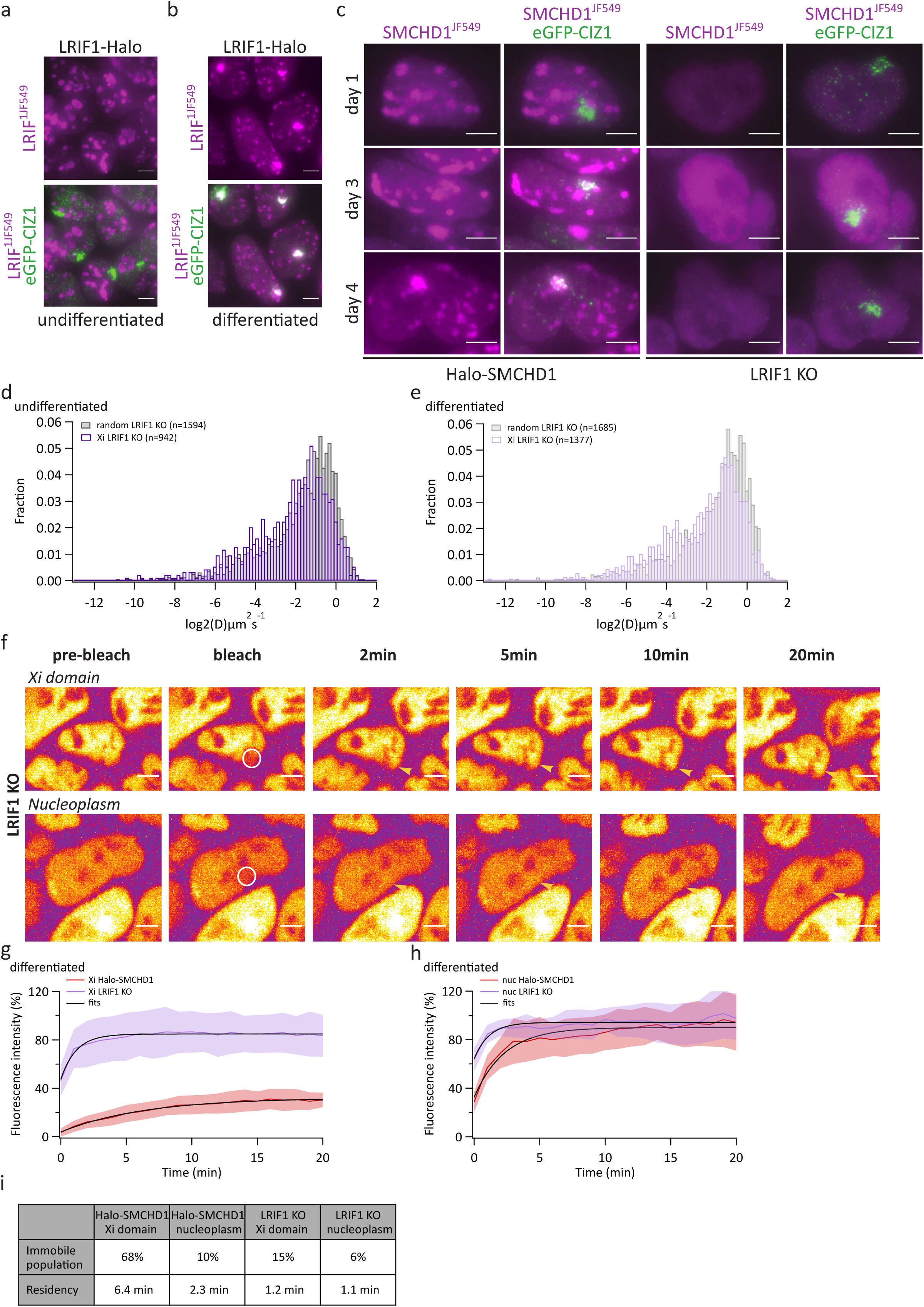
Absence of LRIF1 leads to loss of the SMCHD1 Xi-specific dynamic behaviour. a-b. Representative live-cell images of LRIF1-Halo mESC line, taken after 24h doxycycline addition (a) or 7 days of differentiation (b). c. Representative live images showing gradual enrichment or no enrichment of SMCHD1 over the Xi domain marked by eGFP-CIZ1 during differentiation in Halo-SMCHD1 and Lrif1^-/-^ mESCs, respectively. All images are maximum-intensity projections, scale bar - 5µm. d. In dark purple, same as in Figure 3d. In dark grey, normalized histogram of diffusion coefficients of tracks from a random region in undifferentiated Lrif1^-/-^ mESCs, kept for 24h with doxycycline. e. In light purple, same as in Figure 3e. In light grey, normalized histogram of diffusion coefficients of tracks from a random region in Lrif1^-/-^ mESCs differentiated for 7 days. f. Representative images from a FRAP experiment using the Lrif1^-/-^ mESCs differentiated for 7 days. The bleached region is marked by a white circle on the bleach snapshot and by a yellow arrow in the post-bleach snapshots. Scale bar -5µm. g-h. Single exponential fits of the average fluorescence recovery curves over the Xi domain (g) or a random nucleoplasmic region (h). i. Summary of the immobile population and residency values extracted from the fits shown in g-h.

**Extended Data Figure 5.**
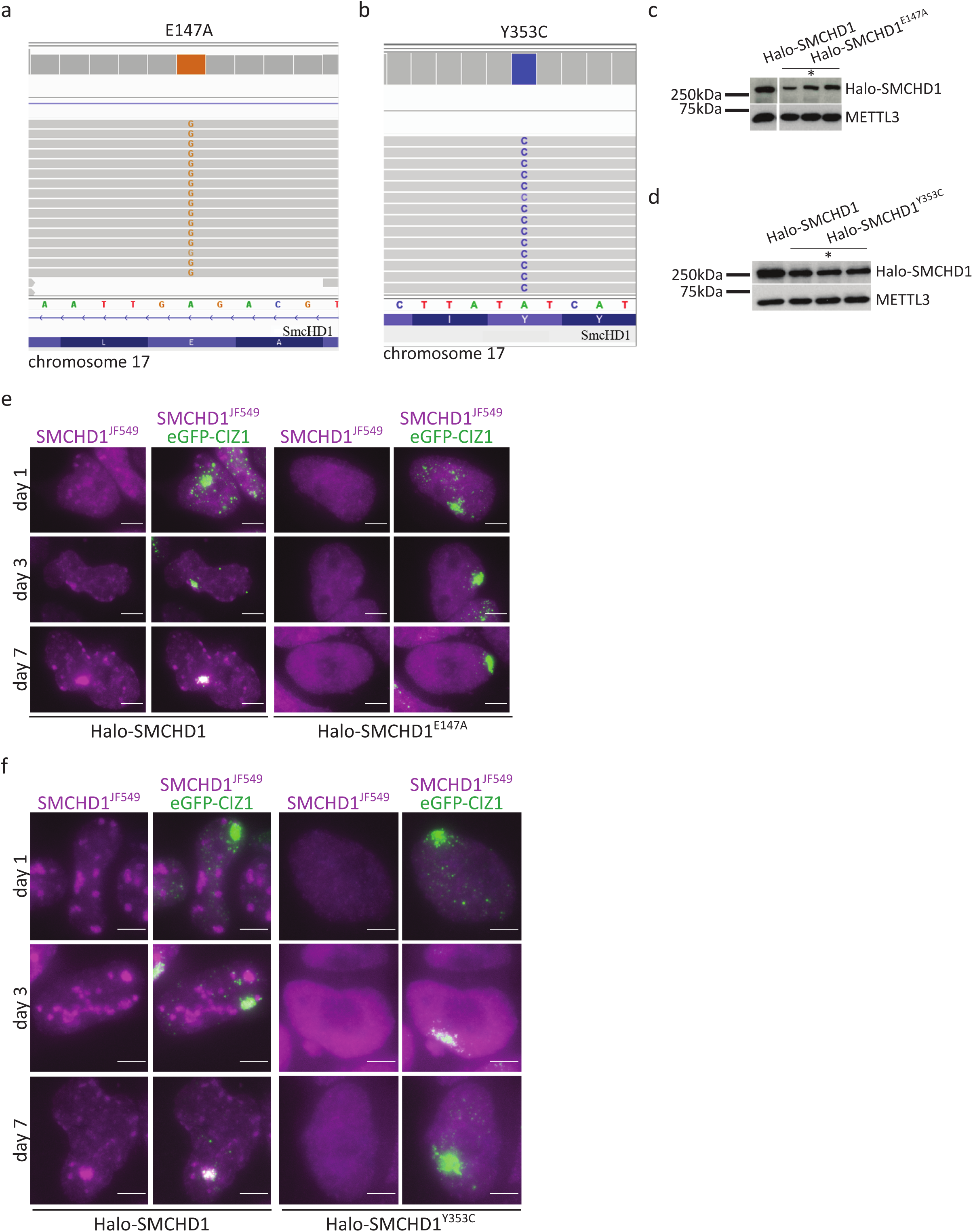
SMCHD1 GHKL ATPase loss-of-function mutations alter the protein’s stability and accumulation over the Xi. a-b. Genome browser screenshot of chrRNA-seq reads showing homozygosity for the selected Halo-SMCHD1^E147A^ (a) or Halo-SMCHD1^Y353C^ (b) clones. c-d. Western Blot showing the expression of SMCHD1 in independent Halo-SMCHD1^E147A^ (c) or Halo-SMCHD1^Y353C^ (d) clones. * marks the clones which were selected for further experiments for each cell line. In c, the extracts were run on the same gel but in non-adjacent lanes. e-f. Representative live images showing there is gradual enrichment or no enrichment of SMCHD1 over the Xi domain marked by eGFP-CIZ1 during differentiation in Halo-SMCHD1, Halo-SMCHD1^E147A^ or Halo-SMCHD1^Y353C^ mESCs. Images are maximum-intensity projections, scale bar - 5µm.

**Extended Data Figure 6.**
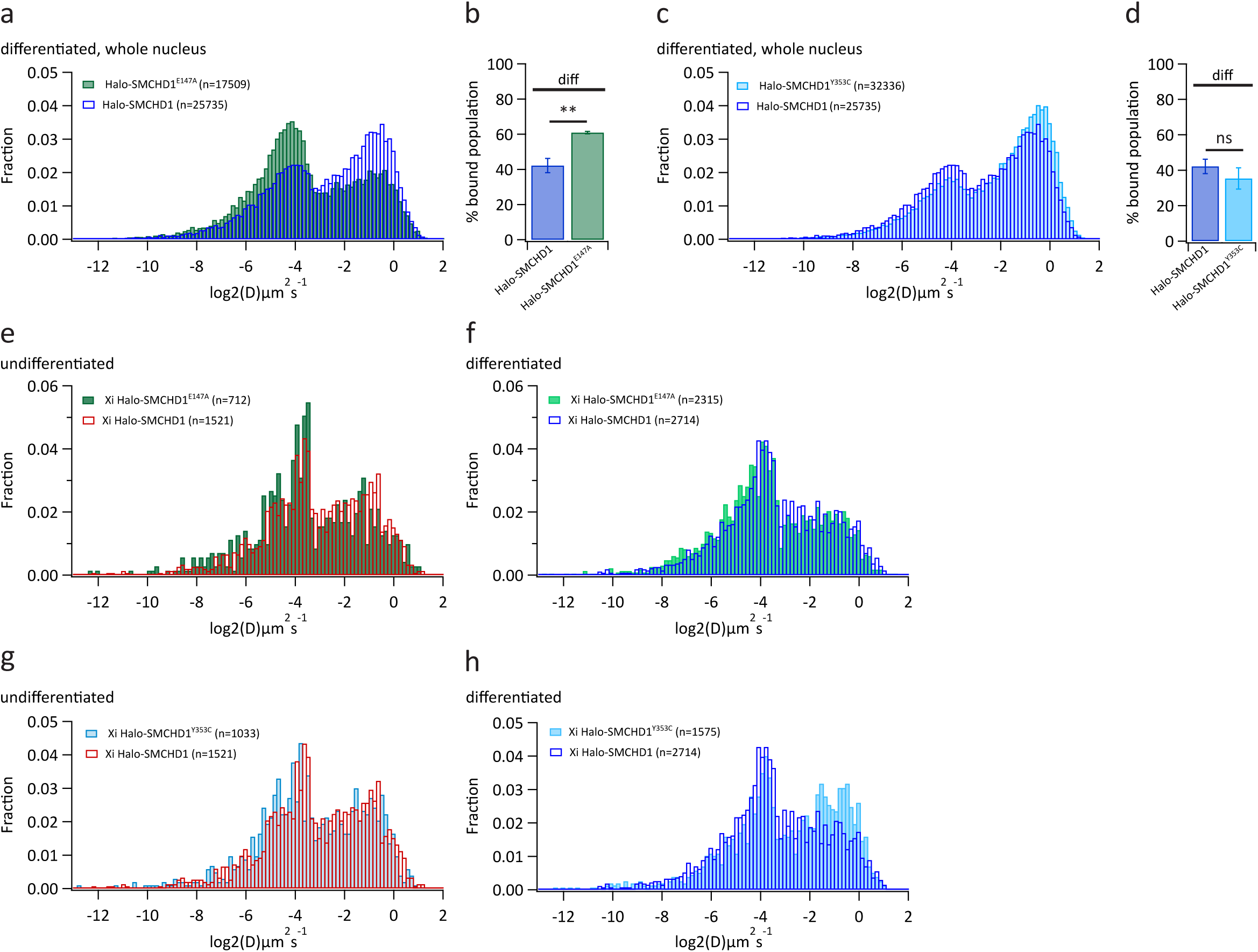
Characterization of the dynamic behaviour of SMCHD1 GHKL ATPase loss-of-function mutants using SPT. a. Normalized histogram of diffusion coefficients of tracks from across the whole nucleus in indicated cell lines differentiated for 7 days. b. Bar plots showing the average percentage of bound molecules of SMCHD1 (3 independent replicates each). Error bars represent the STDEV (unpaired t-test). c. In blue, same as in a. In turquoise, normalized histogram of diffusion coefficients of tracks from across the whole nucleus in Halo-SMCHD1^Y353C^ mESCs differentiated for 7 days. d. Bar plots showing the average percentage of bound molecules of SMCHD1 (3 independent replicates each). Error bars represent the STDEV (unpaired t-test). e. In red, same as in Figure 3a. In green, same as in Figure 5e. f. In blue, same as in Figure 3b. In light green, same as in Figure 5f. g. In red, same as in Figure 3a. In turquoise, same as in Figure 5h. h. In blue, same as in Figure 3b. In light blue, same as in Figure 5i. diff = differentiated

**Extended Data Figure 7.**
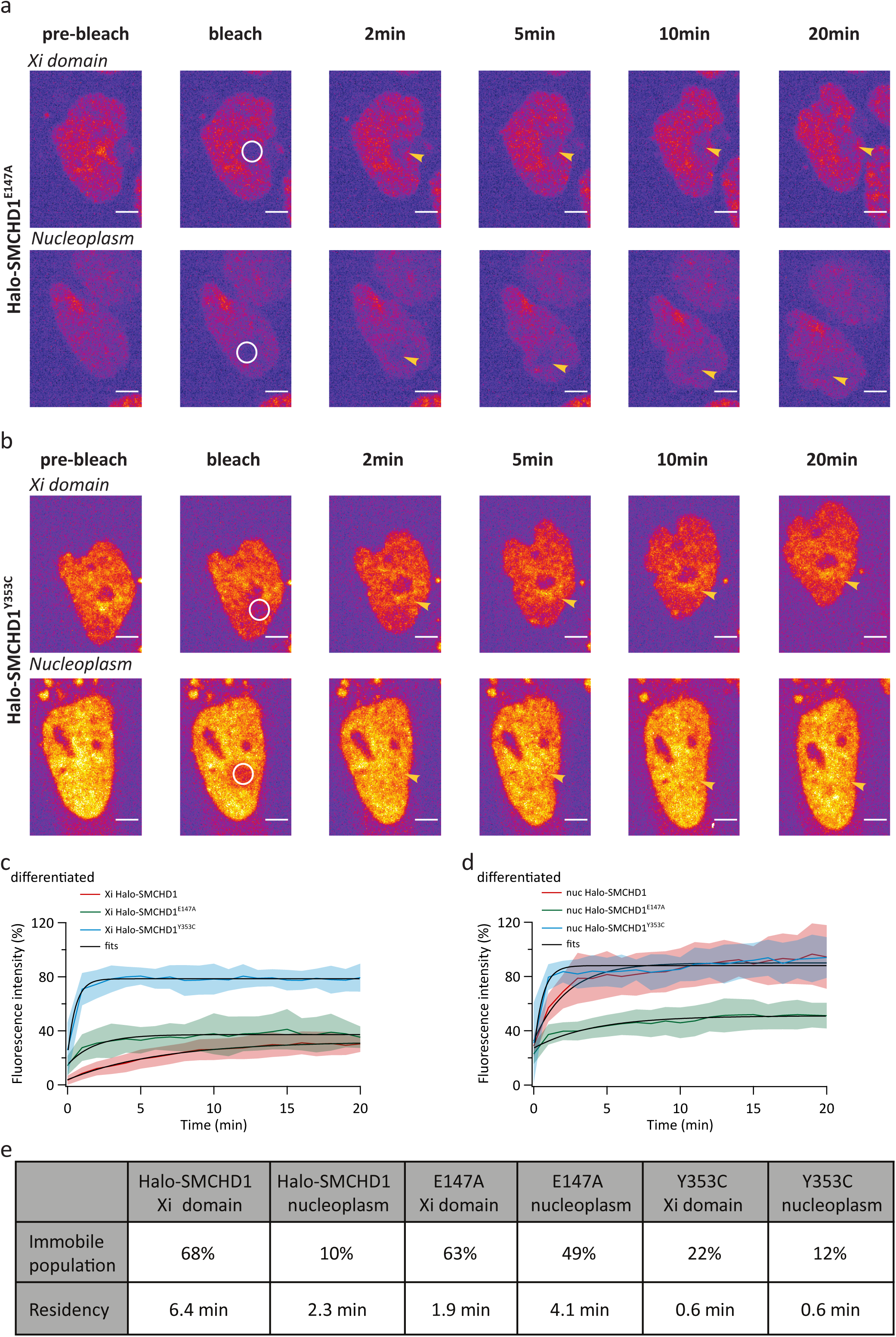
FRAP analysis of the dynamic behaviour of SMCHD1 GHKL ATPase loss-of-function mutants. a-b. Representative images from a FRAP experiment using Halo-SMCHD1^E147A^ (a) or Halo-SMCHD1^Y353C^ (b) mESCs differentiated for 7 days. The bleached region is marked by a white circle on the bleach snapshot and by a yellow arrow in the post-bleach snapshots. Scale bar - 5µm. c-d. Single exponential fits of the average fluorescence recovery curves over the Xi (c) or a random nucleoplasmic region (d). e. Summary of the immobile population and residency values extracted from the fits shown in c-d.

**Extended Data Figure 8.**
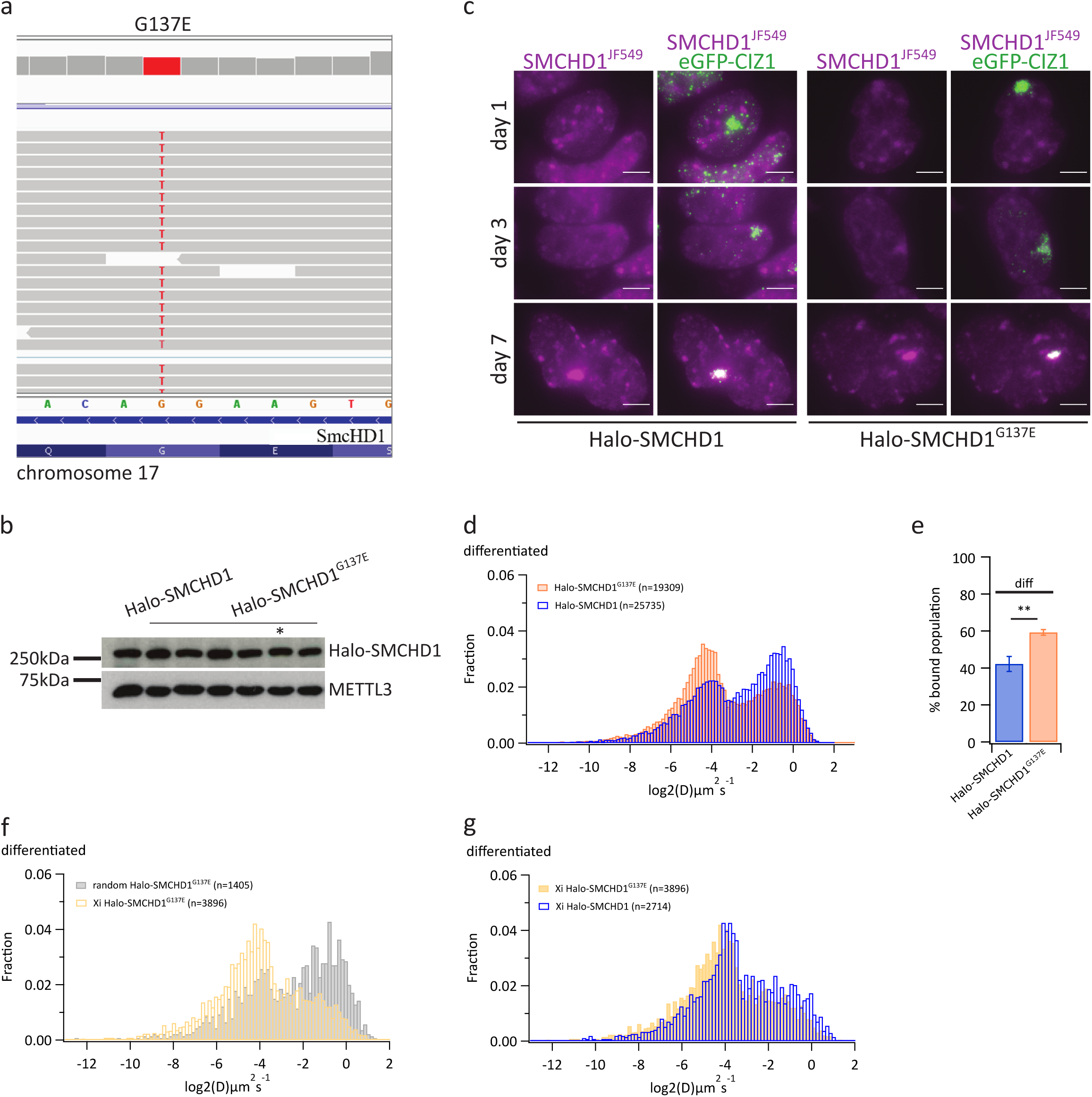
Characterisation of effects on a GHKL ATPase gain-of-function mutation of SMCHD1 on its stability, Xi association and SPT-determined dynamic behaviour. a. Genome browser screenshot of chrRNA-seq reads showing homozygosity for the selected Halo-SMCHD1^G137E^ clone. b. Western Blot showing the expression of SMCHD1 in Halo-SMCHD1^G137E^ clones. * marks the clone selected for further experiments. c. Representative live-cell images showing gradual enrichment of SMCHD1 over the Xi marked by eGFP-CIZ1 during differentiation in Halo-SMCHD1 and Halo-SMCHD1^G137E^ mESCs. Images are maximum-intensity projections, scale bar - 5µm. d. In blue, same as in Extended Data 6a. In orange, normalized histogram of diffusion coefficients of tracks from across the whole nucleus in Halo-SMCHD1^G137E^ mESCs differentiated for 7 days. e. Bar plots showing the average percentage of bound molecules of SMCHD1 (3 independent replicates each). Error bars represent the STDEV (unpaired t-test). f. Normalized histogram of diffusion coefficients of tracks from across the Xi (yellow) or a random region (light grey) in Halo-SMCHD1^G137E^ mESCs differentiated for 7 days. g. In blue, same as in Figure 3b. In yellow, same as in f. diff = differentiated

**Extended Data Figure 9.**
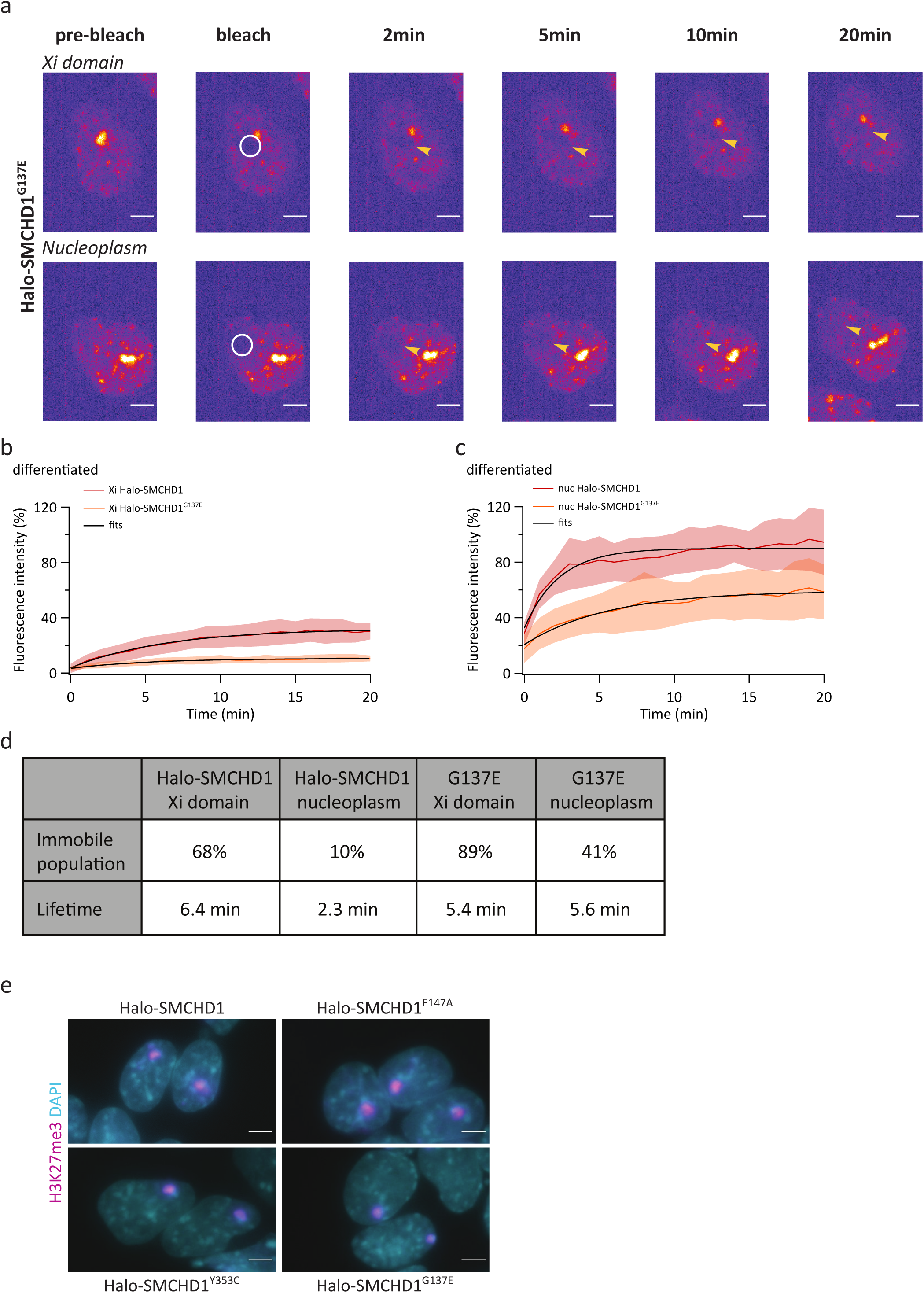
FRAP analysis of the dynamic behaviour of an SMCHD1 GHKL ATPase gain-of-function mutant. a. Representative images from a FRAP experiment using Halo-SMCHD1^G137E^ mESCs differentiated for 7 days. The bleached region is marked by a white circle on the bleach snapshot and by a yellow arrow in the post-bleach snapshots. Scale bar - 5µm. b-c. Single exponential fits of the average fluorescence recovery curves over the Xi domain (b) or a random nucleoplasmic region (c). d. Summary of the immobile population and residency values extracted from the fits shown in b-c. e. Immunostaining for H3K27me3 in Halo-SMCHD1, Halo-SMCHD1^E147A^, Halo-SMCHD1^Y353C^ and Halo-SMCHD1^G137E^ mESCs differentiated for 7 days. Images are maximum-intensity projections, scale bar - 5µm.

